# De novo peptide sequencing with InstaNovo: Accurate, database-free peptide identification for large scale proteomics experiments

**DOI:** 10.1101/2023.08.30.555055

**Authors:** Kevin Eloff, Konstantinos Kalogeropoulos, Oliver Morell, Amandla Mabona, Jakob Berg Jespersen, Wesley Williams, Sam P. B. van Beljouw, Marcin Skwark, Andreas Hougaard Laustsen, Stan J. J. Brouns, Anne Ljungers, Erwin M. Schoof, Jeroen Van Goey, Ulrich auf dem Keller, Karim Beguir, Nicolas Lopez Carranza, Timothy P. Jenkins

**Affiliations:** InstaDeep Ltd, 5 Merchant Square, London, W2 1AY, UK; Department of Biotechnology and Biomedicine, Technical University of Denmark, Kongens Lyngby, Denmark; Novo Nordisk Foundation Center for Biosustainability, Technical University of Denmark, Kongens Lyngby, Denmark; Department of Bionanoscience, Delft University of Technology, 2629 HZ Delft, Netherlands; Kavli Institute of Nanoscience, 2629 HZ Delft, Netherlands

## Abstract

Bottom-up mass spectrometry-based proteomics is challenged by the task of identifying the peptide that generates a tandem mass spectrum. Traditional methods that rely on known peptide sequence databases are limited and may not be applicable in certain contexts. *De novo* peptide sequencing, which assigns peptide sequences to the spectra without prior information, is valuable for various biological applications; yet, due to a lack of accuracy, it remains challenging to apply this approach in many situations. Here, we introduce InstaNovo, a transformer neural network with the ability to translate fragment ion peaks into the sequence of amino acids that make up the studied peptide(s). The model was trained on 28 million labelled spectra matched to 742k human peptides from the ProteomeTools project. We demonstrate that InstaNovo outperforms current state-of-the-art methods on benchmark datasets and showcase its utility in several applications. Building upon human intuition, we also introduce InstaNovo+, a multinomial diffusion model that further improves performance by iterative refinement of predicted sequences. Using these models, we could *de novo* sequence antibody-based therapeutics with unprecedented coverage, discover novel peptides, and detect unreported organisms in different datasets, thereby expanding the scope and detection rate of proteomics searches. Finally, we could experimentally validate tryptic and non-tryptic peptides with targeted proteomics, demonstrating the fidelity of our predictions. Our models unlock a plethora of opportunities across different scientific domains, such as direct protein sequencing, immunopeptidomics, and exploration of the dark proteome.

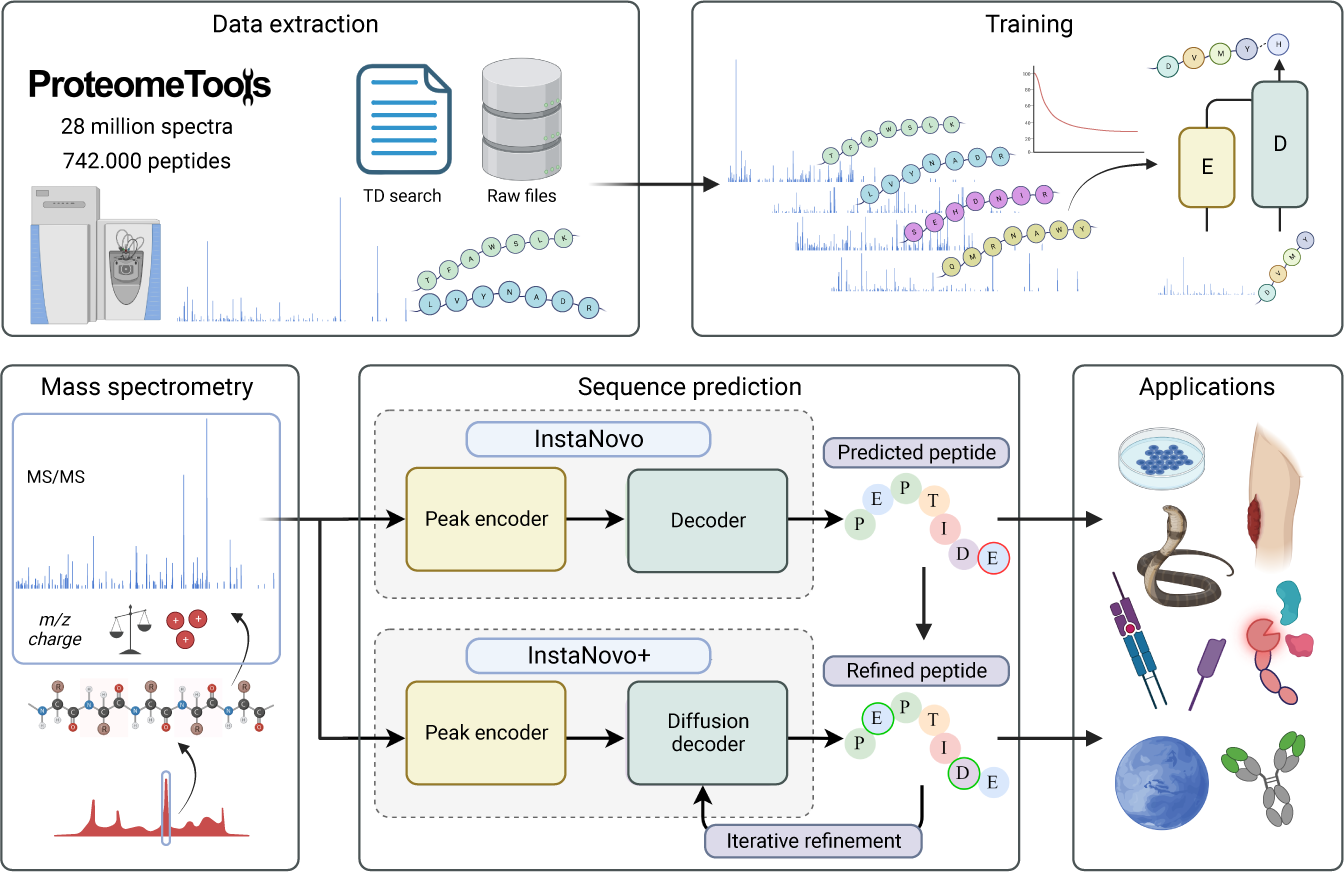

## 1 Main

Mass spectrometry (MS)-based proteomics has revolutionised the way we study proteins on a large scale [1]. Bottom-up proteomics, the main workflow used for system-wide proteomics experiments, relies on the identification of peptides by comparing recorded tandem mass (MS/MS) spectra containing fragment ions, to theoretical peptide fragmentation spectra generated from *in silico* digestion of a protein database [2–5]. Presently, the strategy of database search with target-decoy (TD) FDR estimation is almost exclusively used for both spectrum-centric and peptide-centric acquisition methods [6, 7]. The database search approach allows for peptide scoring against acquired spectra and calculation of false discovery rates (FDR) of the resulting peptide-spectrum matches (PSMs), which are also strictly controlled at the peptide and protein grouping level [8–11]. Although database search with TD FDR estimation presents a convenient and proven way to reduce the computational search space and control FDR in MS-based proteomics, this approach has critical shortcomings [12, 13]. Naturally, a database search narrows the scope of the recorded raw data, and only yields identifications for protein sequences present in the supplied database. Therefore, the selection of the employed database is of great importance, and a poor choice of database can hinder identification of protein isoforms, alternative splicing events, coding single-nucleotide polymorphisms (SNPs), or elucidation of proteins from other organisms not considered for database inclusion. Similarly, database search cannot identify engineered sequences or evolved proteins of interest without knowledge of their sequence and are agnostic to transcription or translation errors. Another major limitation of database search is the skyrocketing cost in search space complexity and its impact on peptide and protein identification. Inclusion of even a relatively modest number of post-translational modifications (PTMs) exponentially increases the computational cost and processing-time of database search [14, 15]. This limits searches to only a few PTMs and makes semi-tryptic or open searches, which would allow for identifications of alternative start sites and proteolytically processed proteoforms, time-consuming and computationally expensive [16, 17]. The expanded search space also results in an increased false positive rate, which causes FDR hikes and therefore lower identification numbers [18, 19].

An alternative approach to database search is *de novo* peptide sequencing, which relies on peptide identification through precursor fragmentation and fragment ion fingerprinting. This approach is the method of choice for bottomup proteomics when prior sequence information is absent [20, 21]. Modern *de novo* sequencing algorithms have attempted to streamline and automate the process of manual fragment identification and peptide sequencing, achieving impressive results [22, 23]. However, such algorithms still suffer from substantial computational costs and high FDRs, rendering *de novo* sequencing for large scale experiments unattainable [24, 25]. Recently, with the advent of deep learning and powerful neural network architectures, as well as the explosion in MS dataset generation and developments in instrumentation, we are experiencing a renaissance in the field of PSM inference [26–29], rescoring and *de novo* sequencing peptide prediction [30–34]. Such approaches hold the promise of accurate peptide identification with linear increases in compute costs for inference, rather than the current exponential cost increases associated with database search. *De novo* approaches represent a powerful methodology for system-wide sequencing experiments without the need for prior sequence information or additional downsides of database search [35]. By overcoming the limitations of database search, *de novo* sequencing opens the door to proteomics applications previously considered out of reach. However, to date such *de novo* sequencing algorithms have not quite met the performance level required to truly leverage *de novo* protein sequencing, and their performance compared to database search remains underwhelming.

Here, we introduce InstaNovo, a model which exceeds state-of-the-art performance on *de novo* peptide prediction with substantial increases in precision and recall rates compared to existing tools. InstaNovo is a transformer model which uses multi-scale sinusoidal embeddings [36] to effectively encode mass spectrometry peaks. These inputs are processed by 9 transformer decoder layers, which cross-attend to the peak embeddings. We apply knapsack beam-search decoding for candidate selection and peptide scoring. We also introduce InstaNovo+, an iterative refinement diffusion model inspired by manual human *de novo* sequencing, which further improves prediction accuracy.

## 2 Results

### 2.1 Training dataset extraction, preprocessing, and InstaNovo model architecture

Consistent with the literature [37, 38], we reasoned that our model architecture would benefit from training with a large, consistent, well documented training dataset. We decided to train our model on the largest available proteomics dataset, which has been recorded with modern, state-of-the-art instrumentation, containing high resolution spectra for peptides of human origin. InstaNovo (IN) was trained on the large scale ProteomeTools [39] dataset, which comprises over 700,000 synthetic tryptic peptides covering the entirety of canonical human proteins and isoforms, as well as encompassing peptides generated from alternative proteases and HLA peptides. We used the data from the first three parts of the ProteomeTools project, and split the database search results into two datasets. The first dataset is derived from the evidence results of the MaxQuant [40] searches available in the repository, and contains the best PSMs per peptide and is therefore referred to as the highconfidence ProteomeTools (HC-PT) dataset. The second dataset contains all PSMs regardless of quality (derived from the ms results of the searches), and is referred to as the all-confidence ProteomeTools (AC-PT) dataset. The HC-PT dataset contains 2.6M unique spectra and the unfiltered AC-PT dataset contains 28M total spectra. Both datasets contain 742k unique peptides (Fig. 1a). Distributions of the dataset properties show expected behaviour in terms of m/z, charge, measurement error, etc. (Supplementary Fig. 1).

**Fig. 1.**
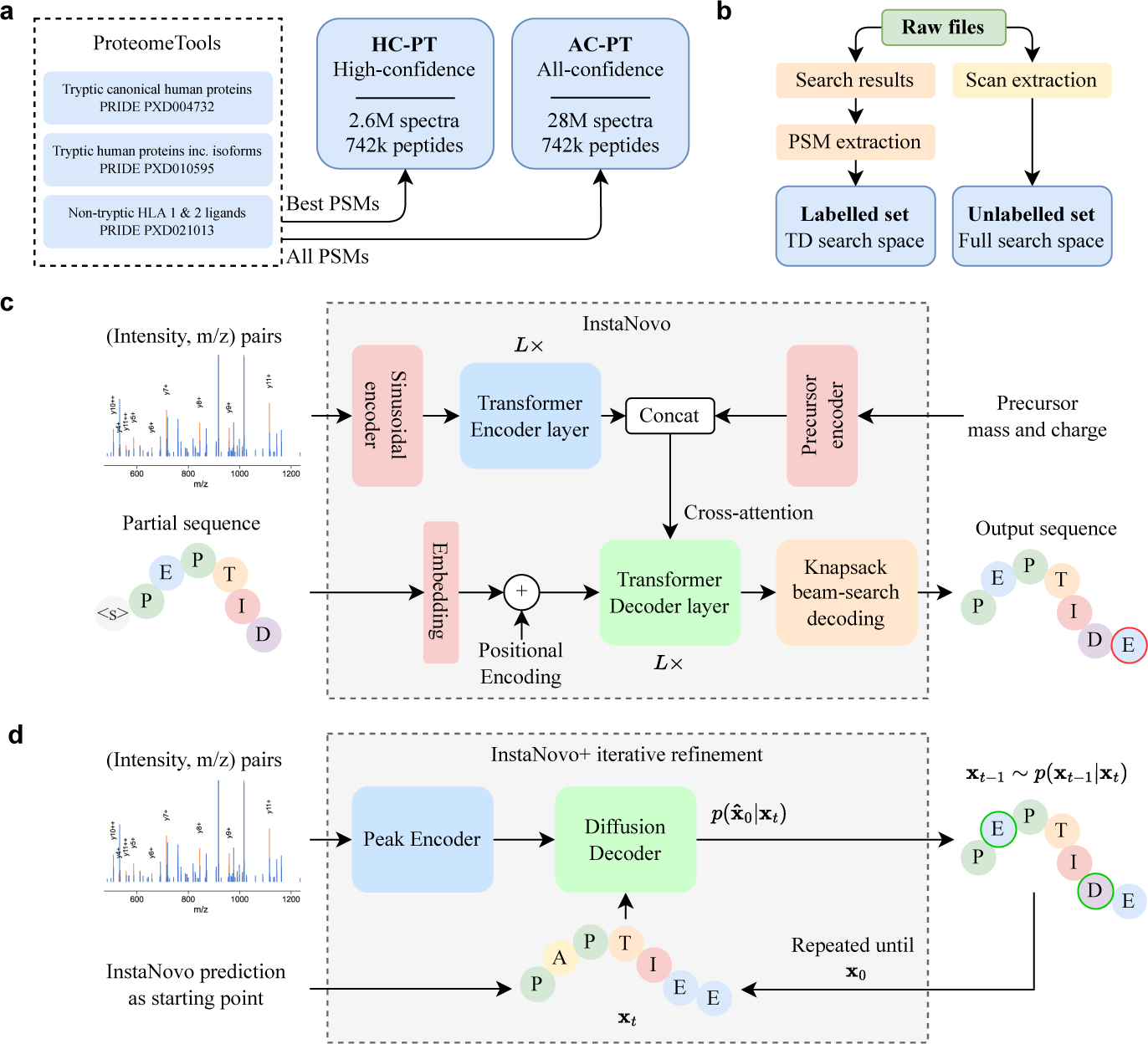
InstaNovo (IN) pipeline overview. **a,** ProteomeTools datasets and their PRIDE repository identifiers. Each dataset covers a unique set of synthetic peptides, derived from human protein sequences, which have been measured with mass spectrometry. **b,** overview of data extraction and preprocessing steps. Raw data were matched with the results of a database search with TD FDR estimation (controlled at 1%) to create the training dataset of our models. **c,** IN model architecture. The model takes a mass spectrum as input, which is transformed to a latent embedding representation using multi-scale sinusoidal embeddings that encodes the intensity and m/z vectors. This is passed through L transformer encoder layers, each with multiple heads to derive a cross-attention representation of the peaks in the spectrum. Additional precursor information is included and concatenated to form the encoder output, which is cross-attended by L decoder layers. The precursor information may alternatively be encoded as the start-of-sequence token in the decoder. The decoder takes in an embedding of the partially decoded peptide sequence, and is responsible for predicting the next residue of the peptide. A knapsack beam-search decoding is applied to ensure the model outputs a confident prediction that matches the precursor mass and charge. **d,** overview over the iterative refinement model, IN+. The model features the IN encoder and a diffusion decoder, which iterates over sequence predictions in a series of timesteps, denoising and refining predictions using a multinomial probability distribution for discrete sequence prediction.

After obtaining the training data from the repository, we devised a pipeline to extract the spectrum information and associated metadata we believed were needed for model training (Fig. 1b and Supplementary Fig. 2). Using available data with HCD fragmentation and pools of 1,000 peptides in individual runs, we extracted the PSM results from database searches, cross-referenced the scan numbers and extracted the spectrum information from the raw data files, obtaining our training input data and labels. We trained our models and previously published tools with both datasets, to assess and compare their performance^1^.

Inspired by recent developments in the *de novo* sequencing field [32, 34], we reasoned that the transformer architecture, ubiquitously recognised as the most uniform and generalisable neural network architecture in machine learning [41–44], would be readily adaptable and applicable for *de novo* peptide sequencing with mass spectrometry data. This is further supported by contemporaneous work [45] that builds on transformer-based *de novo* sequencing models, though there are other architectures that have also shown promising results [46]. The main reason for this is its ability to self-attend and cross-attend to the inputs coming from the spectrum intensity landscape; this enables them to pay attention to the peptide ions that are co-fragmented in the MS2 spectrum and capture their relationships, co-dependence, and distribution to arrive to meaningful sequence predictions. Therefore, we designed our neural network to take the mass spectrum embeddings as model inputs, encoding the intensities and their positions (m/z in the mass spectrum) in the fragmentation spectra. Recent research has shown that mass spectra vectors can be better represented with sinusoidal embeddings [36]. In these encodings, the m/z peaks are encoded with varying frequencies along the hidden dimension of the encoded output. These encodings are then processed with two dense layers, concatenated with the intensity vector, and another two dense layers – providing a high quality encoding of the spectral information. We have confirmed that this approach matches and surpasses other embedding approaches. Hence, we have included this method of latent representation of mass spectra in our model. The encoder has 9 layers, each with 16 heads, a hidden dimension of 768, and a feedforward dimension of 1,024. This encoder allows the fragment ions and their intensities to self-attend to other ions present in the spectra. We also concatenate the MS2 spectrum information with the precursor mass, m/z and charge, allowing the model to account for the precursor information that is useful for prediction, and would be readily available during data acquisition. This precursor may alternatively be encoded as the start-of-sequence token in the decoder, but we found no difference to model performance. The transformer decoder, also consisting of 9 layers with 16 heads each, cross-attends over the encoded spectra. This enables the model to take in the previous residues from the predicted sequence and auto-regressively predict the next token. This simple yet powerful architecture, which is essentially a sequence-to-sequence model, allows for robust learning and next token prediction of peptide sequences from mass spectra. To augment our autoregressive model, we implement knapsack based beam-search decoding. This ensures the model always outputs a peptide sequence that matches m/z of the precursor. This eliminates the need for multiple predictions and retains performance while increasing model confidence and decreasing FDRs in the full search space. IN recall is marginally reduced across datasets (0.05-0.2%) compared to standard beam search with 5 predictions/spectrum, and peptide inference takes longer compared to beam search, but reductions in almost all error types justify its use. Together, this architecture constitutes our IN model (Fig. 1c; Supplementary Fig. 3).

#### 2.1.1 Iterative refinement of predictions improves performance

After our initial model training and promising results in sequence decoding, we speculated that next token prediction is not the most optimal approach to mass spectrum sequence decoding. Since the model is auto-regressive, there is no typical approach to correcting errors made early on during decoding. This is particularly problematic due to the peptide fragmentation properties. The first fragmentation products of a peptide might not even be present in the spectrum due to the first mass cutoff in the spectrum acquisition, or they might not behave ideally in the mass spectrometer due to their physicochemical properties. Beam-search would ideally remedy this, but as the sequence length increases, the issue remains.

Under HCD and CID fragmentation, the most intense ions are the b and y ions [47–50] of the peptide, with the y ions of tryptic peptides generally having better readout properties, potentially due to charge localisation. For that reason, many *de novo* sequencing models start token prediction from the right hand side of the sequence, and that is also what we’re doing for our base model IN. However, we argued that since internal y or even b ions are more intense, and there might be an advantage in exploring approaches that decode the peptide sequence all at once instead of performing next token prediction (see Supplementary Fig. 8). Similarly, we thought that starting residue prediction or sequence order might be wrong, especially in cases where fragment ions are not that intense or spectrum landscapes are noisy. In such cases, the model does not have a chance to update its prediction, or take into account multiple series of ions.

With recent literature showing diffusion models outperforming previous architectures [51–54], we reasoned that probabilistic denoising models would be well suited for our spectrum to sequence prediction. In addition, we believed that the iterative refinement properties of denoising models match well with the way humans approach the problem of *de novo* sequencing, operating with an initial fuzzy prediction based on distinct, unambiguous elements of the spectrum, revisiting and refining the prediction in serial timesteps from previous steps and information from the spectrum in relation to the updated predicted sequence. Based on previous experience [55], we adapted the denoising principles to suit our purpose, and introduced an iterative refinement model that takes an initial prediction (either random or from the IN model), refines and improves on it by revisiting the information encoded by the spectrum given the updated knowledge provided by the peptide sequence. The model, similar in architecture to IN, consists of a transformer encoder and decoder. The decoder, rather than performing autoregressive predictions, iteratively refines predictions in 20 steps. The encoding portion of the model is identical to IN, while the decoder has a fixed sequence length and predicts the full denoised sequence in each model pass. The decoder also cross-attends to an embedding of the current timestep, giving the model an indication on how far along the refinement is.

We termed this iterative refinement *de novo* sequencing model InstaNovo+ (IN+; Fig. 1d and Supplementary Fig. 4). When the IN predictions were used as the starting input sequences to IN+, we saw a considerable improvement in model performance and recall in our validation sets. This indicates that IN+ is adept in recognising errors in the initial predictions and correcting them through refinement of the predicted sequences in a series of steps.

### 2.2 Comparative performance evaluation

We conducted performance evaluation of IN by comparing it to the current state-of-the-art models, including PointNovo [56], and Casanovo [32]. The selection of these models was based on their availability, their representation of comparable architectures and their previously reported performance [31, 57]. We utilised two benchmark datasets, the high-resolution nine-species dataset [33] which serves as a standard benchmark for evaluating deep learning *de novo* peptide sequencing tools, and the ProteomeTools [39] dataset which provides a more comprehensive collection of high-quality mass spectra derived from synthetic peptides. When we assessed the peptide-level precision-recall curve comparing the models trained only on nine-species, and those trained on HC-PT and fine-tuned on nine-species, we see IN+ and IN outperforming Casanovo when fine-tuned, while Casanovo is slightly better when trained from scratch (Fig. 2a). We then trained Casanovo, IN and IN+ trained on HC-PT and evaluated on HC-PT and AC-PT respectively (Fig. 2b and c). On the ninespecies dataset, we evaluated the model accuracy in three setups (Fig. 2d and e). Firstly, the models were trained on nine-species excluding yeast, and then evaluated on yeast. In this setup, the dataset was too small to effectively train IN+, which is why it was excluded from this comparison. Secondly, we compared the models trained on HC-PT and evaluated directly on yeast. In this case, we found IN+ performed the best, outperforming Casanovo, validating our choice of architecture. PointNovo was excluded from these evaluations as it never converged to a comparable level of performance when trained on HC-PT. Finally, we compared the models trained on HC-PT and then fine-tuned on the nine-species exc. yeast. All three models improved significantly, with IN+ reaching a peptide-level accuracy of 54.6%. On HC-PT, the precision-recall curve of IN demonstrated improved calibration compared to IN+, with higher peptide precision for the same recall values. We expect this is due to the way we estimated the lower bound of the diffusion model confidence, which is not as straight-forward as auto-regressive models. In all peptide sets, we find IN significantly outperforming Casanovo, with IN+ marginally improving the IN predictions. We found that though IN+ in itself marginally improves recall, it ends up predicting not only many of the same peptides as IN, but also different ones. As such IN+ does not merely constitute a refinement in our base model, but can be used in addition to IN, overall substantially increasing the number of peptides predicted with low FDR. We use the database search results to ground our search and derive a surrogate confidence threshold for FDR estimation. Comparing the PSMs identified in database search with model predictions, we calculate the confidence threshold of the *de novo* peptide sequencing models that can yield the predictions with 5% FDR. We evaluate the predictions above this confidence threshold that are identical with the database search PSMs. In the nine-species dataset, a database search identified 27,142 PSMs after filtering of a maximum peptide length of 30 and a maximum of 800 peaks in the spectrum. Within that PSM pool, we found that Casanovo predicted 7,580 PSMs at 5% FDR with 446 not found in either IN or IN+; IN predicted 9,244 PSMs (726 unique) which constitutes 21.95% more than Casanovo, and IN+ identifies 10,995 PSMs (2,553 unique), 45.05% more than Casanovo. Together IN and IN+ identified 12,161 PSMs, 60.43% more than Casanovo, which constituted a substantially improved performance of both models individually, as well as when combined (Fig. 2f and g). This trend still held true for the other two datasets (HC-PT and AC-PT), though the improvement was smallest for HC-PT (Supplementary Fig. 5a and b). Error analysis indicated IN and IN+ are incorrectly classifying predictions in the same categories as Casanovo, with IN+ expectedly having increased sequence reordering errors in other datasets (Supplementary Fig. 6). We also assessed the accuracies of each model on HC-PT and AC-PT, grouped by tryptic, HLA-I, HLA-II, LysN, and AspN peptides (Supplementary Fig. 7a and b).

**Fig. 2.**
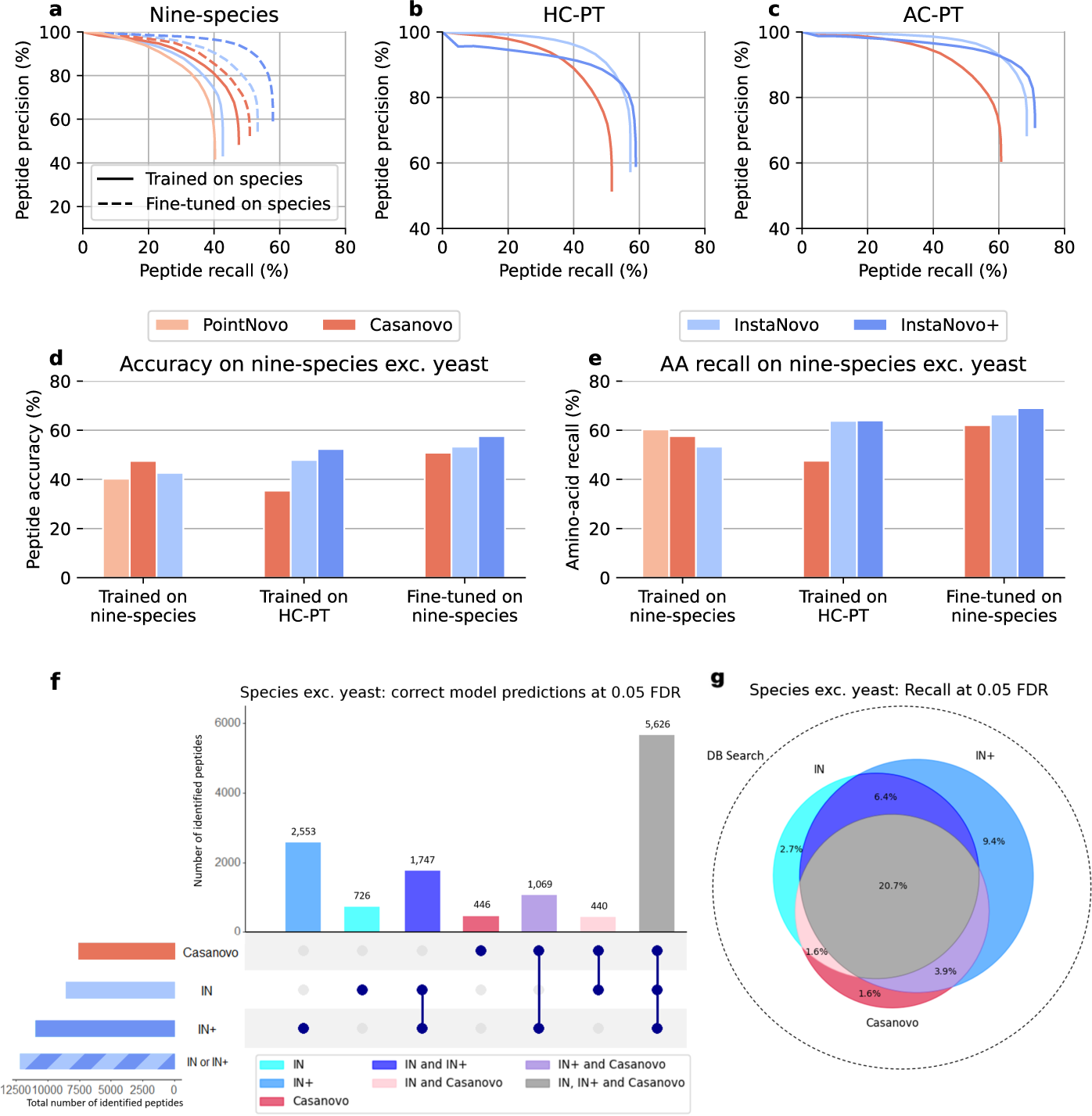
Comparative evaluation of PointNovo, Casanovo, InstaNovo, and InstaNovo+. **a,** Peptide-level precision recall curves on nine-species exc. yeast. **b,** peptide-level precision recall curves on high-confidence ProteomeTools (HC-PT). **c,** peptide-level precision recall curves on all-confidence ProteomeTools (AC-PT). **d,** Peptide-level accuracy of each model on the high-resolution nine-species dataset excluding yeast. The first set of accuracies are trained on nine species exc. yeast, and evaluated on yeast. The second set is trained on HC-PT and then evaluated on yeast. The third set is trained on HC-PT, finetuned on nine species, and then evaluated on yeast. **e,** Amino-acid level accuracy of each model on the high-resolution nine-species dataset excluding yeast. **f,** Peptide-level UpSet plot illustrating the intersection of correct predictions made by the finetuned IN, IN+, and Casanovo models on the nine-species exc. yeast dataset, when evaluated at a false discovery rate (FDR) of 0.05. **g,** Peptide-level Venn Diagram illustrating the same intersections as figure f, but displaying them as percentages (recall) of the DB search ground truth (ms_ninespecies_benchmark) dataset, which is illustrated by the area of the circle with the dotted edge. Areas in the Venn diagram are approximate, due to the imperfection of the Venn algorithm.

### 2.3 Evaluation of InstaNovo on application-focused datasets

#### 2.3.1 InstaNovo shows robust performance and adds value to bottom-up proteomics searches

We evaluate IN and IN+ on 8 external validation datasets. Database search was applied to each, with the search results and number of spectra outlined in Supplementary Table 1. In a given dataset IN achieved up to 72.4% peptide accuracy and IN+ achieved up to 73.6% peptide accuracy (*S. brodae* proteome) without further fine tuning on individual datasets, and only including the training evaluation rounds. However, the performance fluctuated depending on the dataset, resulting in an average of 48.3% peptide accuracy ±19.4% std for IN, and 51.5% peptide accuracy ±21.1% std for IN+ on 8 biological application oriented datasets (Fig. 3a and Supplementary Table 2). At 5% FDR, IN predicts a median of 4,014 PSMs (Fig. 3b), or an average of 34% novel PSMs at 5% FDR compared to the total PSMs in database search results (Fig. 3c). Within the database search results, IN+ finds on average 3% more PSMs which were not covered by InstaNovo, while improving peptide accuracy by 1.5% on average (Supplementary Table 3). Precision recall curves in application-focused datasets show considerable variance depending on sample type and origin (Fig. 3d and e), while model precision as a function of confidence are generally conserved, especially for confidence values above 95%, with the exception of the snake venom proteomics and the nanobodies dataset (Fig. 3f). The IN architecture predicts a peptide sequence with an average 142 ms / spectrum, or 120 minutes for a typical single shot MS run of 50 k spectra. IN+ processes the same number of spectra in 26 minutes, with batch sizes of 64 spectra processed in 1.94s (performance measured on a consumer-level hardware: Intel i5-13600k CPU, RTX 3090ti GPU). When using InstaNovo predictions as a starting point for InstaNovo+, the total runtime would be the sum of the two model runtimes. The model runtime scales linearly with the number of spectra, which is invariant in terms of protein digestion, closed or open searches, type of fragmentation or resolution, or number of modifications (Supplementary Fig. 9). These numbers compare well with search times for conventional proteomics analysis platforms with database search implementations. We expect clear improvement in runtime and PSM detection rates in datasets searched against large protein databases or datasets with a large amount of spectra. We observed in multiple instances that our models predict sequences from spectra with high confidence, while database searches match the same spectra to different sequences. In some of those cases, analysis with spectrum similarity correlation metrics indicates that our model predictions match the experimental spectrum better than the database search results (Supplementary Fig. 10). This suggests that the model can improve the precision of database search PSMs, if applied in the context of an auxiliary or re-scoring search engine in database searches. We monitor predictions of novel peptides with targeted proteomics experiments, validating our model experimentally (Supplementary Fig. 11). These results demonstrate that the model predicts bona fide peptide sequences without any prior information, and can inherently discern between good and bad predictions with high fidelity based on the prediction model confidence.

**Fig. 3.**
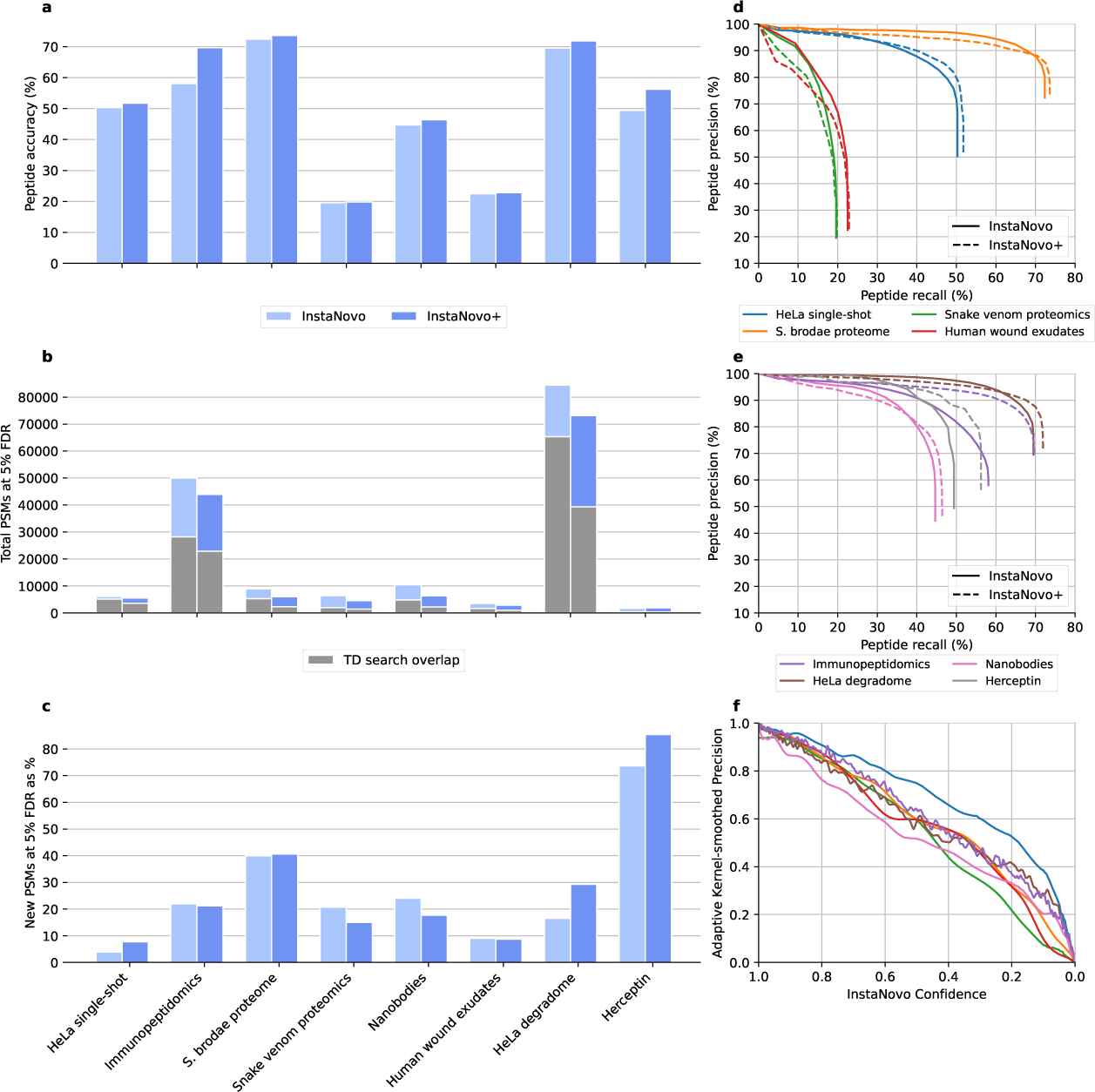
Performance of InstaNovo (IN) and InstaNovo+ (IN+) on the labelled application-focused datasets. **a,** Peptide-level accuracy of IN and IN+ on each application-focused dataset. **b,** Total number of PSMs for IN and IN+ models at 5% FDR. Overlap with database search PSMs is shown in grey. **c,** Novel PSMs at 5% FDR for IN and IN+, expressed as a percentage of database search total PSMs. **d,** Peptide level precision-recall curves for proteomes explored in this study. These consist of HeLa cell lysate proteome, *S. brodae* proteome from a co-enrichment culture, snake venom proteomes, and the proteome from human patient wound exudates as extracted from dressings. **e,** Comparison of peptide-level precision-recall curves for both models on the datasets where novel sequences were involved. These were HLA peptide enriched samples, nanobodies and the antibody Herceptin, as well as a HeLa proteome dataset including semi-tryptic and open search peptides. **f,** Kernel-smoothed precision of model confidence distributions across multiple datasets for IN.

#### 2.3.2 InstaNovo increases PSM rate in HeLa proteomes

First, we wanted to investigate the performance of IN in one of the gold standards in proteomics experiments, the lysate of HeLa cells. HeLa proteomes are frequently used for benchmark studies, and are also routinely used as quality controls between sample batches during data acquisition to assess instrument performance and robustness. We analysed 200 ng of HeLa cells with an Orbitrap Exploris 480 mass spectrometer. IN was able to achieve 49.6% recall in the HeLa single-shot dataset, assigning correct (identical to the database search) sequences for 8,774 PSMs. At 99.9% confidence, we observe 1% false positives, as assessed by grounding to the database search. At 98.1%, we observe 5% false positive hits, indicating that model confidence sufficiently captures prediction certainty, and suggesting optimal model thresholds that might be used for relevant FDR cutoffs. Using the same confidence cutoff (5% FDR) for sequence predictions, IN increased the database search PSM identification rate by 7.5%, identifying 1,338 more PSMs in the MS/MS scans that did not result in any database search hits. Using IN+, we detected 365 more correct PSMs from database searches, even though the total PSM rate did not improve for this dataset. Using the average of output amino acid confidences for each prediction, generating 5 predictions for each spectrum and filtering for sequences that match the precursor mass, we observed in most cases (n=11,979, 87,4% of subset where any prediction matches the precursor mass, 67,74% of the complete PSM space) the first prediction matches the precursor mass, suggesting that the model outputs the correct sequence with the prediction possessing the highest confidence score, where possible (Fig. 4a). Next, we wondered whether our additional predicted sequences would match to the human proteome without any prior database information, and whether the protein coverage and PSM rate would be increased. By choosing a cutoff of 98% prediction confidence, equating to 5% FDR for IN (Fig. 4b) and performing similar thresholding for IN+ (Fig. 4c), we could assess model predictions at a low FDR rate, as per standard practice in proteomics experiments. Considering only IN predictions that match the precursor mass, we were able to naively match 4,029 PSMs corresponding to 3,836 unique peptide sequences, mapping to 1,595 proteins in the human database. It is worth noting that prediction of longer sequences that map to a standard reference proteome, even at lower confidence thresholds, is a direct way of obtaining matches with low FDRs. The chances of obtaining a hit at random with a peptide sequence of 6 residues, usually the lowest peptide length allowed in mass spectrometry, is approximately 1 in 1,600. When considering a length of 10 residues (the mode of peptide length observed experimentally), the number approaches 1 in 4 billion of getting a hit by chance. These approximations were also what we observed in simulations with randomly generated databases of equal size to the human proteome and the same prediction confidence threshold, where the few random matches encountered were 6-8 residues long (Supplementary Fig. 12a), different in length distribution compared to the predicted sequences, and significantly lower than the PSM rate observed in the human database (two-sided one sample t-test, p-value=3.9*e*^−26^). The few random matches to these generated databases from our model predictions were obtained for database search PSMs at the lower end of the search engine’s score and posterior error probability, validating once again our model and its confidence as a suitable metric for FDR estimation. For our predicted human proteome matches, we could observe that the mode of the peptides per protein distribution is 1, and follow a hypergeometric distribution, similar to PSMs or peptides per protein distribution of database searches in bottom-up proteomics (Supplementary Fig. 12b and c). Similarly, the peptide length distribution is in agreement with the database search results. IN also predicted peptide sequences with 1 or more missed cleavages, in the same relative amount as database searches, indicating an adequate performance on peptides that contain internal lysine or arginine residues (Supplementary Fig. 12d). Satisfied with the performance of our base model, we evaluated our knapsack algorithm for selection of model outputs. With a 74% confidence threshold in our knapsack predictions, which are multiplied amino acid probabilities across the peptide sequence (corresponding to 5% FDR), we detected 4,527 PSMs mapping to 4,308 unique peptide sequences and 1,729 proteins. Given the superior performance of our knapsack predictions, we henceforth evaluated the models and report results with our knapsack implementation. When analysing the full search space, without any restrictions to the database search results, we obtained similar prediction confidence, peptide length and protein coverage distribution. However, by utilising the full search space (all MS2 spectra), and using identical confidence filters, we are able to retrieve 6,105 predicted PSMs (Fig. 4d), mapping to 5,287 unique peptides and 2,214 proteins in our protein database. Among these, we observed 811 more unique peptides and 151 more proteins that match to the human proteome, and would otherwise go undetected by database searches (Supplementary Fig. 12e and f). These numbers correspond to a 29.09% and 20.01% detection rate increase on peptide and protein level, respectively, comparing to the restricted database search space predictions when applying these filters, and a 3.84% and 3.63% peptide and protein identification increase respectively when compared directly to database search results, with no post-prediction filtering and with strict FDR thresholds. We observed gains and new peptide matches in new proteins as well as proteins with previously sequenced peptides, increasing coverage at all levels. Importantly, we noticed several examples of peptides in the database search space where the model predicted sequence theoretical fragmentation spectrum, whether binary or Prosit predicted, correlated better with the experimental spectrum than the database search PSM sequence (Fig. 4e). Using the same cutoffs, we also obtained 442 PSMs with 419 unique peptide sequences that do not map to any human protein. We used protein Blast to determine the origin of those sequences, with the highest confidence sequences corresponding to lysyl endopeptidase, which was used in digestion of the HeLa proteome along with trypsin. Other sequences map to proteins in the human proteome with a single mismatch of asparagine to glutamate residue, a common modification in proteomics experiments. These results suggest that IN generates high confidence predictions which support and expand database search results even in the most comprehensively characterised proteomes.

**Fig. 4.**
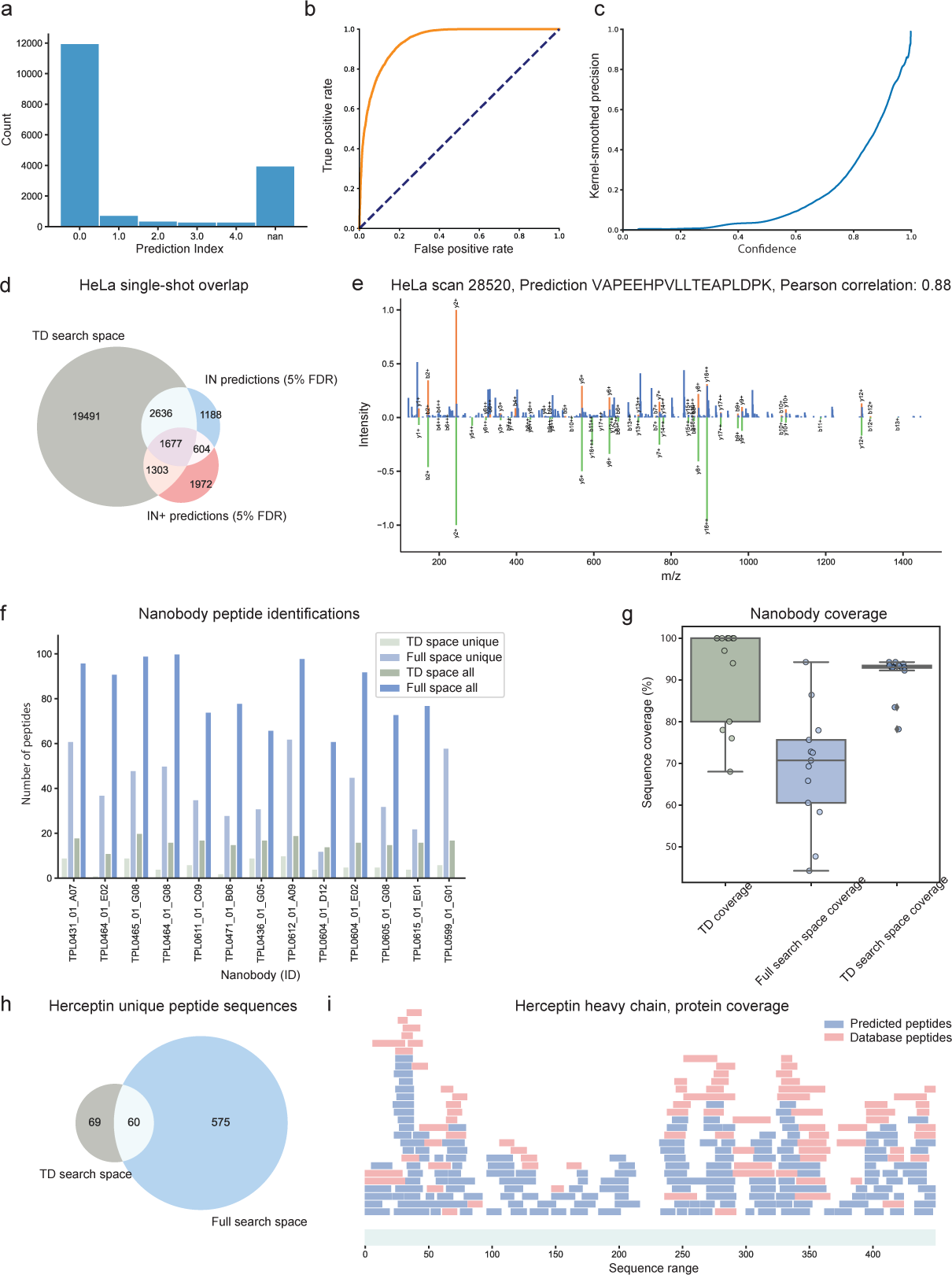
InstaNovo (IN) achieves good accuracy on the established HeLa proteome, and sequences therapeutics in different formats. **a,** Barplot of prediction distribution index with the highest confidence matching the precursor mass. **b,** ROC curve analysis for HeLa single-shot proteome IN predictions. **c,** IN+ prediction confidence in the HeLa single-shot proteome. **d,** IN and IN+ predictions and their overlap with database search PSMs at 5% FDR in the HeLa single-shot proteome. **e,** Mirror plot of experimental spectrum (top) and Prosit predicted spectrum (bottom), in a prediction sequence exhibiting better correlation than the database search PSM. **f,** Barplot of total and unique peptides for the nanobodies analysed. **g,** Sequencing coverage for nanobodies analysed for database search, IN predicted database search, and IN predicted full search at 5% FDR. **h,** Venn diagram for peptides sequences matching to herceptin in the 6 protease digests analysed with database search and IN predicted in the full search space. **i,** PSMs for database search results and IN predicted peptides for the herceptin heavy chain.

#### 2.3.3 InstaNovo can sequence engineered biomolecules with near total coverage in a single run

Next, we wanted to investigate our model’s performance in *de novo* sequencing of novel, engineered biomolecules. We expressed and purified 13 nanobody scaffolds raised against different snake toxin antigens with phage display technology, sequenced them and analysed them with mass spectrometry, using a standard sample preparation workflow with trypsin digestion. As the nanobodies were expressed in E. coli, we ran our database search with the E. coli reference proteome and the 13 nanobody sequences as our background database. In our database search, we detected all 13 nanobodies in our samples with 4,465 PSMs, and sequenced them with 91% average protein coverage, 7 of which with 100% sequence coverage. Out of all proteins detected, we achieved 44.15% peptide recall with IN when querying the database PSMs. When applying IN to the sequencing of our nanobodies with 92% confidence threshold (expected FDR of 5%) in our database search space, we obtained an average protein coverage of 68.93% (13.46% standard deviation), with a median of 16 peptides per protein, and 5 unique peptides per nanobody. When evaluating IN on the database search PSM associated scans, we find 4,955 PSM predictions that match our nanobody sequences with 94 unique peptides (Fig. 4f). This increased detection rate could be mapped to an average of 91 (standard deviation of 13.7) peptides per nanobody, improving our coverage to 91.39% (4.66% standard deviation), reflecting a near 23% increase in sequence coverage compared to the database search space (Fig. 4g). Notably, we obtained 7,536 matches mapping to 613 peptides when expanding the search to the full search space (all MS/MS spectra) of our runs, a 6 fold peptide detection increase compared to the PSM space from database searches (Fig. 4h). The unique peptide sequences detected for a nanobody increased to 40, a striking 8 fold increase in average unique sequences when contrasted to the database search space. These surprising results can be for the most part attributed to semi-tryptic or non-tryptic peptides, products of aminopeptidase activity or degradation, which are distinguished by the ragging patterns of their sequences that differ by one terminal residue at a time. These peptides would be missed in traditional database searches, which limit the computational search space by only considering fully tryptic peptides for the theoretical digest and peptide spectrum scoring. Among the novel peptides discovered in the full search space, there were four peptides mapping to a region of a nanobody with ambiguous genomic sequencing results (C09), elucidating the sequence of that region and demonstrating its promise in similar applications (Supplementary Fig. 13). IN predicted 5,068 novel PSMs at the same confidence threshold in the full search space, a 24.83% increase in PSM detection rate compared to the database search. IN+ detected 2,016 novel PSMs at 5% FDR, whilst slightly increasing the overall peptide recall. We also applied our model to a publicly available dataset evaluating mass spectrometry based antibody sequencing [58], where the authors used 9 different proteases and 2 fragmentation activation types to sequence herceptin (commercially available as Trastuzumab), a monoclonal humanised antibody used to treat breast and stomach cancer by binding to the HER2 receptor [59, 60]. The reason for this was to evaluate our model in a different antibody format prediction, as well as to assess model performance in prediction of peptides generated with several different proteases and fragmentation schemes. Combining the database searches for a subset of 6 out of the 9 different proteases, we detected 1,796 PSMs mapping to 129 unique peptides in the heavy and light chain of herceptin, covering 63.02% of the heavy chain with 83 peptides, and 71.96% of the light chain with 46 peptides. IN achieved 68.99% peptide recall, while it expands detected sequences by 575 unique peptide sequences at 5% FDR, obtaining similar detection rate increases to our nanobody sequencing results. Importantly, it increases protein coverage to 92.87% and 100% for heavy and light chains, respectively (Fig. 4i). Interestingly, IN assigns the correct sequence for PSMs generated by all proteases tested (27.63% PSM recall), with lowest success rates for LysN (12.28%) and highest for thermolysin (56.45%). Surprisingly, IN predicts correct sequences for a fraction of MS2 scans obtained with EThcD fragmentation, even though no such scans were included in training of the model, albeit with a lower PSM recall rate (17.93% across all 6 proteases). These results indicate that our models are adept at novel protein sequencing, matching database results coverages while eliminating several steps in the workflows, as the sequencing could be performed directly on the protein level without necessitating prior genomic information. IN is capable of prediction of peptide sequences of various origins, and has a surprising, however limited, success when predicting sequences from similar fragmentation schemes (Supplementary Fig. 14). This has the potential to substantially speed up novel therapeutics sequencing, by cutting down on sample preparation time, increasing robustness, and decreasing points of failure. Furthermore, by measuring protein and their levels directly, there is considerable promise in assessment of binding efficiency from protein binder abundance.

### 2.4 InstaNovo detects pathogens in human patient wound fluids

Following these results, we questioned how our model would perform in complex samples where the presence of multiple organisms is suspected. For that, we utilised wound fluid exudates from human venous leg ulcer patients, analysed in a previous study [61]. These chronic wounds are prone to infection, therefore we suspected that we could detect potential pathogens present in the wound exudates, which would be missed when analysing the raw data with the human proteome alone. In addition, these samples possess plasma-like complexity with a high protein dynamic range, posing another challenge we would like to assess our performance on. Our model achieves 22.24% peptide recall when tested against the database searches. This is the second lowest peptide recall observed across all datasets tested, and we speculate that this is due to the high dynamic range of wound exudate proteins, and experimental noise in MS/MS scans. With IN, we correctly predicted 849 out of 3,727 database PSMs that could be mapped to the human proteome, belonging to 609 unique peptide sequences and 407 proteins. As expected, the protein best mapped is human serum albumin (ALBU_HUMAN, P02768) with 124 PSMs and 24 unique peptides. By expanding our search to the full MS/MS space from the two runs corresponding to two different wound dressings applied to the wound at different timepoints, IN predicts 2,981 novel PSMs at 5% FDR, a 14% PSM and 14.8% unique peptide detection rate increase compared to database search. The resulting predictions map to 1,804 unique peptides and 624 proteins, rivalling database search results. IN+ detected 1,307 novel PSMs at 5% FDR, whilst slightly decreasing the overall peptide recall. In this dataset, we observed several expected correct predictions under the 5% FDR cutoff, with 10,980 matched PSMs to 5,794 peptides from the same human database without any FDR thresholds. We extended albumin mapping to 1,225 PSMs with 254 unique peptides (most semi- or non-tryptic), a 10 fold increase compared to the database search space, and observed analogous results in other proteins (Fig. 5a). Notably, these peptides increased albumin sequence coverage from 35.63% in our predictions from the database search space, to 85.22% in the full search space (although falling short of the database search PSMs coverage which was 92.93%). To confirm our hypothesis, we extended our proteome database to include 4 more reference proteomes from pathogens commonly found in wound fluids (*E. Coli, P. aeruginosa, S. aureus* and *Citrobacter sp.* [62–64]). A small fraction of IN at 5% FDR were sequences mapping to pathogens of interest. Even without FDR thresholds, the expected value of randomly mapped hits to the pathogens with a database of that size would be approximately 40, with probability of false positives at 6-8 residues, different from the results we obtain, indicating that some of these hits are bona fide PSMs. We mapped unique sequences to 5 of *P. aeruginosa*, 23 of *E. Coli*, and 24 of *Citrobacter* sp. proteins, with a significant number of sequences mapping to multiple proteomes. Conducting a search with the same database containing the pathogen proteomes, confirmed the presence of the same pathogens in our samples with 80 protein and 130 peptide groups, which demonstrates a robust performance of our models in detecting additional proteins originating from unknown species in biological samples.

**Fig. 5.**
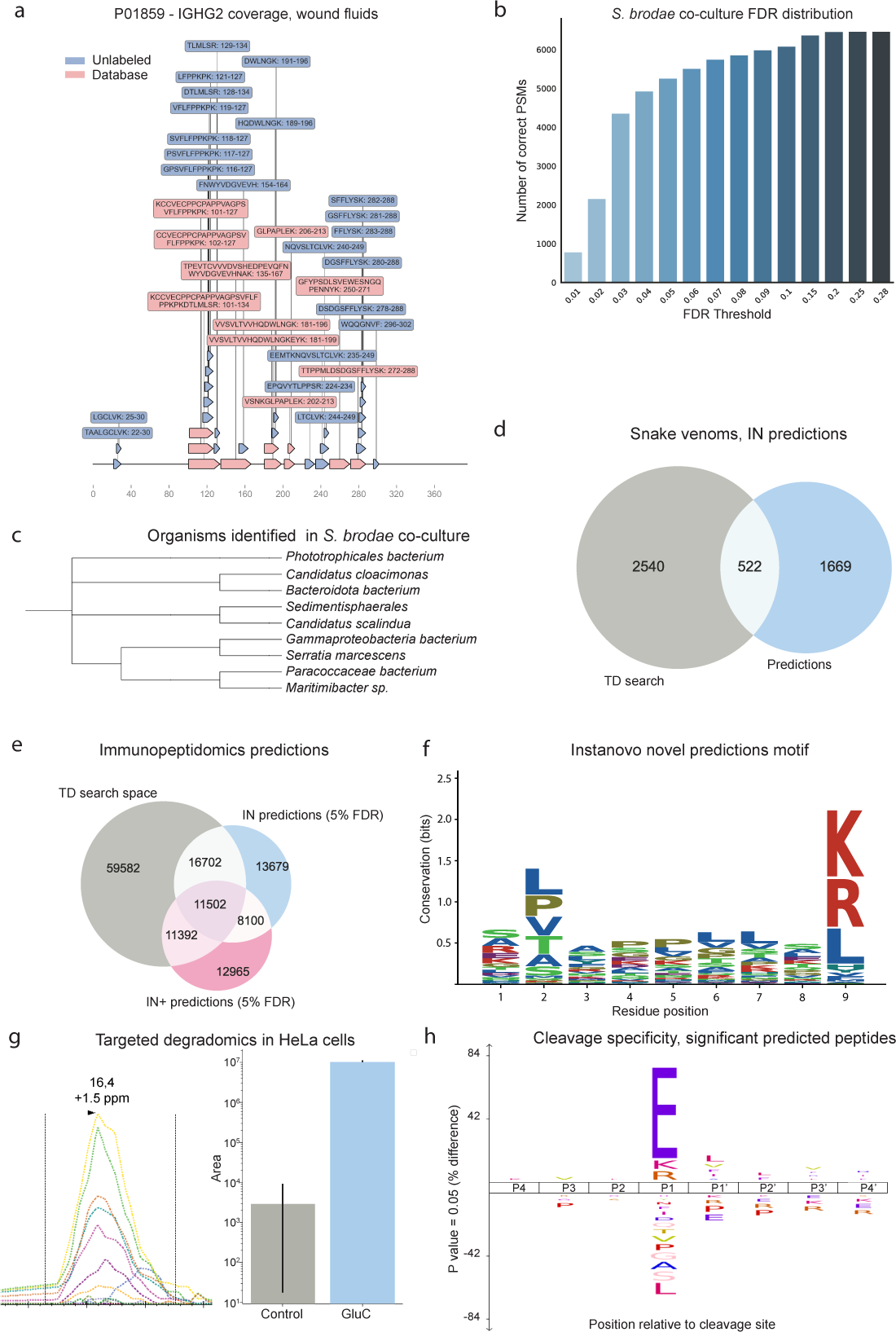
InstaNovo (IN) increases protein coverage, identifies novel organisms, and detects semi- and non-tryptic peptides. **a,** Protein coverage and peptide sequences for P01859 - IGHG2 in human wound fluids, where database search peptides and novel predictions with IN are shown. **b,** Correct PSMs in different precision thresholds in the S. brodae proteome. **c,** Phylogenetic tree of a representative sample of additional organisms identified in the S. brodae culture. **d,** Venn diagram of database search and novel IN predictions of peptide sequences at 5% FDR from snake venom proteomics that map to the proteomes database used. **e,** Venn diagram of database search, IN and IN+ predictions at 5% FDR peptide sequences matching the proteome database used from immunopeptidomics dataset. **f,** Shannon information content of residues in sequence positions of immunopeptidomics experiments. **g,** PRM monitoring of fully GluC-generated peptide ATVWIHGDNEENKE, and its abundance in the two conditions. **h,** GluC specificity profile from statistically significant predicted PSMs matching database search results.

#### 2.4.1 InstaNovo identifies additional organisms in complex bacterial communities

Building upon our previous results, we next questioned how IN performs in the field of metaproteomics, i.e. in complex samples where multiple organisms are present. We chose a co-culture of an enrichment reactor for the marine bacterium *“Candidatus Scalindua brodae”*, which cannot be grown in isolation as of present, and therefore are not classified as a species yet. We hypothesised that due to its inability for self sustained growth, we would detect other organisms present in the culture. IN achieved a recall of 71.5% at full coverage when evaluated against database search results, the highest observed ostensibly due to low proteome complexity. At 1% FDR, we recorded an 8.77% recall, reaching 58.2% at 5% FDR, indicating a need for post-processing tools that sharpen confidence and widen the distance between true positives and false positive results for use in < 1% FDR searches (Fig. 5b). In addition, IN predicts 3,076 more PSMs in the full search space at 5% FDR, a 33.98% increase compared to database search results. Using IN+ and 5% FDR, we detected an additional 293 novel PSMs from database searches and observed a small increase in the total recall, while predicting 2,192 novel PSMs. In the database search space of 9,053 PSMs satisfying our input criteria, IN correctly predicted 5,402 PSMs mapping to 4,896 unique peptide sequences and 1,460 proteins at 5% FDR. When predicting sequences for all MS/MS scans and the same FDR equivalent of model confidence, we match 6,701 PSMs that map to proteins in our *S. brodae* database, corresponding to 6,126 unique peptide sequences and 1,531 proteins, of which 3,330 peptide sequences are novel, while extending protein identifications by 26. At the same FDR, we predicted 2,106 high confidence predictions that remained unaccounted for when searching against the *S. brodae* proteome. When checking against a database comprising sequences for some of the organisms identified with metagenomics in this culture63 and reference proteomes of related *S. brodae* species, we identified 88 predictions matching 82 peptides from 74 protein sequences of several additional species (*S. rubra*, *“Candidatus Kuenenia stuttgartiensis”*, *Geobacter sp.*, and *Sulfurovum sp.*), 41 of which are sequences with 9 residues or longer. We examined the rest of the 1,937 sequences that do not map to any of our databases by comparing them to sequences in genome databases. Using pBLAST and filtering with low expected values (<0.0001) and high identities revealed potential additional species present in our samples, such as *Phototrophicales bacterium, Candidatus Scalindua arabica, Phycisphaerales bacterium, Bacteroidota bacterium,* and *Gemmatimonadota bacterium* (Fig. 5c). These results demonstrate IN is suitable for metaproteomics applications, where multiple organisms are present in analysed samples, with no prior knowledge about presence of these organisms required.

### 2.5 InstaNovo reveals novel peptides in snake venom and enables comprehensive venomics profiling

Next, we wondered if we could apply our models to samples where limited genomic information is available, and there is a potential for novel sequences to be discovered and new insights to be gained. We therefore picked a dataset that recently described the proteome composition of 26 medically relevant snake venoms from sub-Saharan Africa [65], arguing that since not all genomes are available and these proteomes were searched against a pan-snake proteome database, we might detect potential novel sequences unique for some of these species. IN model achieved recall of 18.85% (potentially due to amino acid preference and divergence from training data). IN+ detected an additional 731 correct PSMs from database searches whilst slightly decreasing the overall peptide recall. However, IN expands on the database search results by predicting 5,565 more PSMs in the full search space at a confidence threshold equivalent to 5% FDR, resulting in 1,669 novel peptides and 303 novel protein identifications from the same protein database. Strikingly, these predictions constituted a 54.5% increase in peptide and 46.19% in protein detection rates with our *de novo* sequencing approach, significantly expanding database search results (Fig. 5d). Out of the high confidence predictions in the full search space, we identified 4,113 PSMs that match our protein database, while 2,284 PSMs and 1,117 unique peptide sequences are novel. Using protein Blast and an expected value of < 0.0001 in the top high confidence, long sequence predictions, we could detect matches with high similarity to existing sequences. For example, “SLGGVTTEDCPDGQNLCFK” aligned with isoform 1 sequence of MTLP-2 from *N. kaouthia*, a snake species which was absent from our input dataset. This situation echoes the common occurrence in venom research, where the closest homology match often stems from a different snake species or region and highlights the limitations of relying solely on existing references. Additional matches like “LHSWVECETGECCDQCR” mapped to a snake venom metalloprotease from *E. ocellatus* or “DQGCLPDWSFHEGHCYK” and “DEDCLPDWSSHEGHCYK” mapped to C-type lectins from *Bitis sp.* with a single substitution. Together, these results indicate that these are novel hits with undetected, or not included in the database, search sequences. These can provide insights into novel proteins, isoforms or SNPs in these samples and aid current development efforts for rationally engineered next-generation antivenoms.

### 2.6 InstaNovo discovers new HLA peptides in immunopeptidome experiments

Subsequently, we asked whether our *de novo* sequencing models could be applied to the sequencing of HLA peptides for the analysis of immunopeptidomics experiments. A considerable fraction of our training dataset consisted of HLA peptides, so we were curious to investigate our performance in such evaluation datasets. We chose to evaluate our models with a published interferon induced immunopeptidomics dataset, which was a part of a larger study in high throughput immunopeptidomics [66]. In the complete dataset with a model confidence equivalent to 5% FDR, IN predicted 40,224 PSMs that could be mapped to 8,860 unique peptides and 5,377 proteins in the same human proteome database used in the study, as well as 3,049 peptides in 9,759 PSMs with no matches to the human proteome. Out of the PSMs identified with IN, 28,204 were common with the TD search results, corresponding to a percentage of 28.43%. Remarkably, IN predicts 3,495 novel peptides compared to the TD search, increasing peptide identification rate by 41.53%. IN+ at 5% FDR detected 11,392 more PSMs from the TD search and predicted 12,965 novel PSMs (Fig. 5e). The predicted peptide length formed a distribution centred around 9 residues, with the vast majority (>95%) of predictions being within 8-11 residues. The 9-mer peptides identified with IN showed a motif consistent with MHC bound peptides, exhibiting preferences for certain residues in positions 2 and 9, supporting the model predictions (Fig. 5f). These results indicate that IN performs well in open searches, is adept in prediction of HLA peptide sequences, and can substantially enhance identification rates in immunopeptidome datasets.

### 2.7 InstaNovo can help detect proteolytic products in terminomics experiments

Finally, we questioned our model’s performance in limited processing or degradomic samples, where proteolytic substrates and their discovery are of interest. We hypothesised that our model would perform well in sequencing of semi- or non-tryptic peptides, as our training dataset contained non-tryptic peptides generated with other proteases, similar to the HLA peptides investigated above and results observed in other datasets. Proteolytic processing is an ubiquitous PTM in health and disease, degrading or activating various proteins dependent on signalling and stimuli from the environment. We applied our model to a HeLa proteome incubated with GluC or without GluC in triplicate, before preparing them for MS analysis with standard workflows. GluC is a protease cleaving substrate at the C-terminal side of glutamate residues, therefore we sought to detect such peptides with high confidence to evaluate IN’s performance protease substrate detection. IN achieved a recall of 68.84% in this degradomic dataset, when assessed against a semi-tryptic database search for control and GluC treated samples. The improved performance in this dataset, which is also a HeLa proteome, indicates that multiple shots of the same proteome or increased number of PSMs results in better model predictions and accuracy. We mapped 66,000 predicted PSMs with a 5% FDR to the human proteome in the database search space, while this number increased by 13.84% (75,140 predicted PSMs) in the full search space. These predictions corresponded to 22,425 unique peptides in full search space, a 20% increase compared to the database search space. When contrasted with the database search results, IN predicted 4,635 new peptide sequences and improved peptide detection rate by 11.29%, and protein detection rate by 7.11% (Supplement Fig. 14a and b). IN+ detected an additional 5,696 correct PSMs from the database search, whilst slightly increasing the total recall. Importantly, IN predicted 1,222 new sequences that match the protease profile, i.e. are preceded by glutamate residue in the respective protein sequences these peptides map to (Supplementary Fig. 15c and d). This reflected a 21.84% increase in putative protease cleavages compared to the database search results. Subsequently, we wondered whether these cleavages reflected bona fide peptide detections that were missed by database searches. We therefore revisited our samples and interrogated them with targeted proteomics and PRM precursor monitoring. We were able to identify several high confidence, semi-tryptic or fully GluC generated peptides, and monitor their fragmentation transitions in both conditions (Fig. 5g). When we matched the IN predicted sequences with the abundance computed with the database search results and performed statistics on them to discern significant changes compared to our controls (two-sided two sample independent t-test, filering for log2 fold change > 2 and p-value < 0.01), we could obtain a specificity profile with glutamate significantly over-represented at P1 position, just before the cleavage site (Fig. 5h). These results confirm our hypothesis that IN can be applied to the detection of protease substrates at a system-wide scale, and underscore potential applications in degradomics experiments with semi-tryptic or open searches. Similarly, we expect our model to be widely applicable in the detection of truncated proteoforms, exopeptidase processing, and protein degradation.

## 3 Discussion

By expanding the scope of proteomic applications and providing insights into previously inaccessible protein landscapes, *de novo* peptide sequencing is a promising tool for advancing our understanding of a wide range of complex biological systems. Here, we introduce the IN and IN+ models and analyse their predictive performance in several application domains, including the sequencing of engineered biomolecules, immunopeptidomics, and exploration of the dark proteome. We demonstrate improvements in peptide searches and computational costs, and benchmark across two other tools used for *de novo* sequencing, i.e. PointNovo and Casanovo. To our knowledge, these results represent a major improvement over state of the art algorithms for *de novo* sequencing in bottom up proteomics and constitute a promising step in replacing or complementing database searches.

Beyond the general improvements over state-of-the-art *de novo* peptide sequencing tools, we present applications of our model in several questions in biology. We uncover novel biological findings across eight different datasets, including the identification of proteins in HeLa cells undetected by database search, the expansion of the immunopeptidomics dataset by 175% more peptides, and the characterisation of novel proteolytic cleavages. Given our results and the diversity of the datasets explored in this study, we expect that the model may generalise with high accuracy and satisfactory performance across organisms and biological samples. We anticipate future applications of the model in several other research areas, such as proteogenomics [67], gut microbiome studies [68], studies aiming to explore unreported proteoforms [69]. We also hope our models find suitable applications in the emerging field of single cell proteomics, where increasing PSM detection rates from minute sample amounts is of paramount importance [70, 71].

We expect that by fine-tuning our models on specific tasks, such as big datasets or individual PTMs, they will learn to recognise novel natural or induced chemical modifications of peptide sequences, expanding its applications in chemoproteomics, PTM detection and discovery, as well as multiplexed proteomics. We also expect our models to generalise well to lower resolution spectra and various fragmentation techniques. However, further research is needed to assess the performance and generalisation of IN and IN+ performance and generalisation in different types of mass spectrometers (e.g. instruments with time-of–flight or ion trap detectors), different resolution of MS/MS scans and their effect in performance and prediction confidence, as well as different fragmentation techniques for PTM discovery. We await investigation of different acquisition schemes, such as DIA, and model input adaptation by the creation of pseudo–MS2 spectra [72, 73], facilitating higher detection rates even for applications requiring very high sensitivity.

Following recent trends [74, 75], we anticipate hybrid searches with multiple orthogonal methods of PSM predictions, downstream rescoring algorithms and ensemble models to be increasingly useful in utilising the full recorded spectrum space and maximise detection rates. It has to be noted that in our characterization and evaluation of the model, we consider database search PSMs as ground truth for peptide detection in our dataset. This assumption might be flawed, as database search space PSMs and confidences might be incorrect or incomplete. We believe our models can efficiently be used to corroborate, correct, and/or disprove database search PSMs, increasing detection rates and improving peptide prediction precision. We also speculate that comprehensive post-processing evaluation of model predictions and multivariate filtering based on peptide features and spectrum similarity will increase sensitivity and fidelity of PSMs. Post-processing filters could also serve as a funnel for refinement of predictions with our IN+ model, further leveraging the iterative refinement of predictions with diffusion, which currently is only scratching the surface of its potential. We further believe our models perform adequately well in prediction of non–tryptic peptides, especially if fine-tuned to allow for the use of different peptidases for proteolysis and thereby increasing protein coverage and sequencing. We predict that deep learning approaches will be critical in overcoming the complexity of database searches, and we expect reduced search times for ultra fast sequence predictions in digestion-agnostic proteomics searches.

Together, our results and those of others showed that scale is the most determining factor in *de novo* peptide sequencing model performance, as with other fields where the transformer architecture was employed [38]. We expect to further increase model performance by taking advantage of the vast amount of MS datasets available in repositories. We also anticipate widespread adoption by peers, and look forward to further exploration of fine tuning, protein inference and assembly, as well as building applications on top of our base model for hybrid or *de novo* searches.

## 4 Methods

### 4.1 Data

#### 4.1.1 Training dataset retrieval and preparation

The datasets used in this study were downloaded from the PRIDE repository. Specifically, Parts I-III of the dataset were selected because they contained all human peptides from canonical and isoform proteins, as well as non-tryptic peptides generated by HLA, LysN, and AspN. To ensure a consistent analysis, only the 3x high-energy collision-induced dissociation (HCD) data were utilised, as they provided an inclusion list and employed 3 different HCD fragmentation energies. The raw data files were converted to mzML format using the Proteowizard MSConvert tool [76], with default settings. The result files obtained from MaxQuant [40] (“evidence.txt” or “msms.txt” for high confidence or full dataset, respectively) were employed to extract scan indices for identified peptides, as well as the associated metadata (precursor mass, charge, measurement error, retention time) for each peptide-spectrum match (PSM). To facilitate further analysis, the pyOpenMS python [77] wrapper of the OpenMS C library was utilised. This tool enabled the reading of mzML files, extraction of scans, and association of the scans with the PSM metadata. To refine the dataset and set a padding threshold for the model input features, PSMs were filtered based on specific criteria. Only peptides with a length of 30 or fewer residues and a maximum of 800 peaks in the spectrum were included in the analysis. The resulting data were organised into individual data frames for each run, and subsequently merged to create a consolidated dataset for model training. The key features incorporated into the dataset included mass values, intensity, “MS/MS m/z,” charge, and modified sequence. The same procedure was followed for testing and validating the model. In cases where additional validation was required on all MS2 spectra recorded for each data acquisition run, a separate validation dataset was created. This dataset included all of the MS2 spectra from the raw files (selecting the 800 most intense peaks from each spectrum if more were present), as well as associated metadata, i.e. charge, precursor mass, retention time, and measurement error.

#### 4.1.2 External dataset retrieval and processing

External datasets were also processed using the same methodology as with the training dataset. Publicly available and in house datasets were used to benchmark and validate the model. Specifically, the antibody dataset [58] available in this study was used for benchmarking against similar *de novo* peptide sequencing approaches. The immunopeptidomic dataset [66] used for validation of the model in HLA peptide performance and efficiency originated from this study, and can be found in the PRIDE with identifier PXD006939. The snake venom dataset was downloaded from this article [78] and can be found in the PRIDE repository with identifier PXD036161. The wound exudates originated from this article [61] and are available in PanoramaWeb with dataset identifier PXD025748. The herceptin dataset was found in a *de novo* sequencing tool comparison study [58] in Figshare ((doi.org/10.6084/m9.figshare.21394143)

#### 4.1.3 Data splits

We do a 80:10:10 train/validation/test split for HC-PT and AC-PT based on the unique peptide sequences. When splitting, we ensure there is no leakage between HC-PT sets and AC-PT sets (i.e., no HC-PT train samples are present in the AC-PT test set, etc.). All models and hyper-parameters were chosen based on their validation set performance. Test set results were only computed when writing up the manuscript and used for the reported figures. All results shown in the manuscript are reported on the test set. For Yeast, we use the splits as defined in DeepNovo and PointNovo.

### 4.2 Model implementations

#### 4.2.1 PointNovo and Casanovo implementations

We compared our model to state-of-the-art deep learning methods that have been shown to have superior performance, have been replicated and their architecture is of interest, these include PointNovo [56] and Casanovo [32]. Deepnovo was the first method to incorporate the encoder-decoder architecture to *de novo* peptide sequencing [33, 34], while PointNovo is its successor from the same authors [56] adopts an order invariant network structure called T-Net for the prediction of higher resolution data. Both DeepNovo and PointNovo pass their peak encodings to an LSTM or an output layer to predict the next amino acid. Casanovo frames the problem as a sequence-to-sequence problem [32] and employs a transformer encoder-decoder framework to process and predict sequences of amino acids. PointNovo and Casanovo were both retrained using their respective official GitHub repositories (github.com/volpato30/DeepNovoV2 for PointNovo, and github.com/Noble-Lab/casanovo for Casanovo). The input files for PointNovo were prepared using the repository’s data preparation scripts. The models were trained for 30 epochs, considering a total of 12 ion types with a maximum peptide length of 60 and peptide mass of 5,000 Da. The maximum number of peaks per spectrum was set at 400. Fixed modifications for each dataset included carbamidomethylation of cysteine (C + 57.02 Da). Variable modifications for all datasets were oxidation of methionine (M + 15.99 Da), deamidation of asparagine (N + 0.98 Da) and glutamine (G + 0.98 Da). It is important to note that only carbamidomethylation and oxidation of methionine were observed in the dataset analysis of the Prosit dataset.

#### 4.2.2 Development of InstaNovo architecture

The IN architecture is also based on the transformer encoder-decoder architecture [79]. Similar to PointNovo [56] and Casanovo [32], we represent our MS2 spectra as the set of *N* peaks (**m, I**), where **m** = *m*_1_*, m*_2_*, . . ., m_N_* and **I** = *I*_1_*, I*_2_*, . . ., I_N_* represent the sets of m/z and intensity respectively. To encode these peaks, we employ multi-scale sinusoidal embeddings [36]. We process these encoded peaks through a transformer encoder layer, allowing the model to self-attend and extract relative information between the peaks. The encoder output is concatenated with a learnt latent spectra and a representation of the encoding of the precursor. The precursor mass *m*_prec_ and charge *c*_prec_ are encoded with a sinusoidal encoding and embedding layer respectively, after which they are summed to represent the precursor embedding.

The transformer decoder makes use of causal auto-regressive decoding – predicting one amino acid at a time. The partially decoded sequence is encoded through an embedding layer and a standard sinusoidal positional encoding is added. The input sequence is automatically prepended with a start-of-sequence token. The decoder cross-attends over the encoder output, latent spectra, and precursor encoding.

Formally, for a sequence of residues *y*_1:M_ InstaNovo assigns the probability

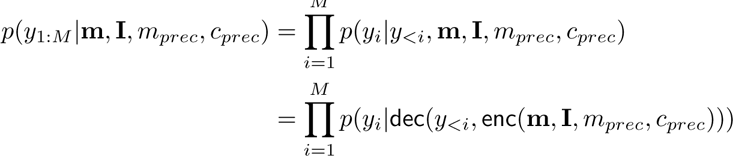

where enc(**m, I***, m_prec_, c_prec_*) are the embeddings returned by feeding the spectra and precursor mass and charge into a transformer encoder as described above and dec(*y_<i_*, x) are the embeddings returned by a transformer decoder fed sequence *y_<i_* and cross-attending on tensor x. *p*(*y_i_|***v**) is parameterised as a linear layer followed by a softmax and we calculate perform training and inference using log-probabilities for numerical stability^2^.

InstaNovo has 95M parameters in total. To train IN, we implement the model in PyTorch [80], with PyTorch Lightning [81] being used to handle the training loop. The loss function computes the cross-entropy between the predicted model logits and the ground truth peptide. All training and model hyperparameters are provided in Supplementary Table 4.

#### 4.2.3 Iterative refinement with InstaNovo+

In addition to IN, we introduce IN+, based on a similar transformer architecture but with a different goal. Rather than auto-regressive decoding, IN+ model is trained to perform multinomial diffusion [55, 82]. This means the model is trained to iteratively remove noise from a corrupted sequence. The full model architecture is given in Supplementary Fig. 4. InstaNovo+ has 170M parameters in total. To perform multinomial diffusion, we define three distributions applied to a discrete fixed-length sequence x*_t_*, where *t* is the current noise step. x_0_ represents an uncorrupted peptide and x*_t_* is a completely corrupted peptide indistinguishable from the starting point x_0_. *T* is a hyperparameter representing the maximum number of noising steps, and is chosen as 20 in this work.

We can now define the diffusion processes. Firstly, the forward noising function *q*(x*_t_ |* x_*t*−1_) defines the distribution over x*_t_* given x_*t*−1_. Secondly, the denoising function *q*(x_*t*−1_ *|* x*_t_*, x_0_) defines the distribution over x_*t*−1_ given x*_t_* and x_0_. Finally the denoising function *p*(x_*t*−1_ *|* x*_t_*) defines the distribution over x_*t*−1_ conditioned on x*_t_* and the model output 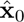:

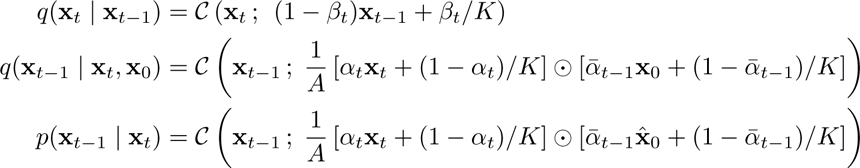

Where *C* denotes a categorical distribution. *β_t_* is the noise schedule, *α_t_* = 1 *− β_t_*, and 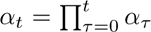 . The noise schedule *α_t_* follows a cosine decay from 1 to 0 as *t* increases [82]. 1/A denotes a normalising constant. The model predicts 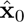 given the spectra, precursor information, the current timestep, and the corrupted sequence x*_t_*. To train the model, we compute the KL-divergence between the true denoising distribution *q*(x_*t*−1_ *|* x*_t_*, x_0_) and the model’s distribution *p*(x_*t*−1_ *|* x*_t_*). Once trained, we can iteratively refine a sequence starting from random noise to arrive at a final prediction for the spectra. Alternatively, we can refine a prediction made by IN, which yielded the best results. To do this, we start from the IN prediction at *t* = 15, and iteratively refine till we reach x_0_. This serves as our IN+ prediction. If the IN+ prediction does not satisfy the precursor mass, we instead fall back to the IN prediction used at *t* = 15. We can also estimate the log probability of IN+ predictions to extract model confidence. We sample a few x*_t_* given the model prediction as x_0_, then take the average KL-divergence loss of the model predicting 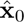 given x*_t_*. This serves as a lower-bound estimate of the log probability. When decoding IN+, we decode 5 samples for each spectra. The sequence that matches the precursor mass with the highest log probability under the model is selected as the IN+ prediction. In the case where we start with an IN prediction and none of the IN+ predictions satisfy the precursor mass, we instead use the IN prediction (which should always fit the precursor).

### 4.3 Metrics and benchmarks

We use peptide recall as our main benchmarking metric for testing and validation datasets. As this is the more stringent of metrics used in *de novo* sequencing algorithm evaluation, we believe this metric reflects our model’s performance the best. We also report peptide precision, as well as amino acid residue precision, recall, and error rates for our training and validation datasets. For each spectrum, a model decodes a peptide with some confidence. For a given confidence threshold, the model predicts peptides for spectra where this confidence is higher than the threshold and for other spectra it predicts nothing. An amino acid in a model-predicted peptide matches one in a gold standard peptide if their masses differ by less than 0.1 Da and the masses of their prefixes differ by less than 0.5 Da. A model-predicted peptide matches a gold standard peptide if they have the same number of amino acids and every amino acid in the model-predicted peptide matches one in the gold standard peptide. Amino acid precision and recall are defined as 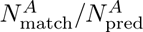 and 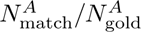 respectively, where 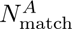 is the total number of matched amino acids in model-predicted peptides, 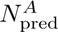 is the total number of amino acids in model-predicted peptides and 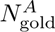 is the total number of amino acids in gold standard peptides. Peptide precision and recall are defined as 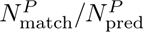 and 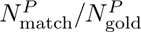 respectively where 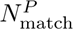 is the total number of matched model-predicted peptides, 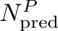 is the total number of model-predicted peptides and 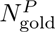 is the total number of gold standard peptides. We further compared our models to baselines using the entire ROC curve rather than just the precision and recall at a single confidence threshold. We obtained these by varying the confidence threshold from the highest to the lowest values obtained in an evaluation dataset and plotting the resulting pairs of (amino acid- or peptide-level) precisions and recall values.

We decoded peptides from our models using beam search with knapsack filtering (Supplementary Algorithm 1). This ensured that the system always found a peptide that fit the precursor mass, improving overall performance and reducing the frequency of almost all individual error types. Beam search (with beam width *B*) is a variant of breadth-first search where at each step, the frontier is pruned to the *B* highest scoring sequences. We use knapsack filtering in beam search to only allow amino acid sequences that can be continued so that their theoretical mass matches the precursor mass to a 50 ppm relative difference. Finding all amino acid sequences whose theoretical mass is in some range is an instance of the knapsack problem and can be solved efficiently using dynamic programming or graph search over an array of possible masses called a *chart*. We precomputed this chart for masses up to 4,000 Da at a resolution of 0.0001 Da using depth-first search (Supplementary Algorithm 2). During beam search, we used the chart to filter out sequences that could not be continued to fit the precursor mass to the tolerance. As a result, all partial sequences on the beam could be continued to fit the precursor mass guaranteeing the system could always find a peptide that matched the precursor to the tolerance.

### 4.4 Application-oriented datasets

#### 4.4.1 Nanobodies

The nanobodies included in this study (Supplementary Table 5) were discovered using phage display technology. Briefly, camelids were immunised with whole venoms from either 8 viperid snake species or 18 elapid snake species, followed by the construction of immune nanobody displaying phage libraries (VIB Nanobody Core, Brussels) as described by Pardon et al. [83] . These libraries were used in phage display selection campaigns including purified biotinylated snake toxins as antigens in a procedure similar to the one described by Ledsgaard et al. [84] with the exception that the nanobody-encoding genes were digested with *Pst1* and *Eco91l* restriction enzymes and subcloned into the xb-145 vector for expression (instead of using *NcoI* and *NotI* restriction enzymes and the pSANG10-3F vector). Individual colonies were picked, cultivated, and used for expression of nanobodies. The periplasmic fractions of the cells (expected to contain the majority of the nanobody protein) were analysed in an expression-normalised capture DELFIA [85]. A subset of nanobodies binding their cognate antigens were Sanger sequenced (Eurofins Genomics) using the M13rev-29 primer. For purification of the nanobodies, *E. coli* BL21(DE) were grown at 37 °C and 220 rpm until OD_600_ reached 0.5, followed by the addition of 0.5 mM IPTG to induce protein expression. After 16 h incubation at 30 °C and 220 rpm, the cells were collected by centrifugation at 4,000 g for 15 minutes at 4 °C and stored at -20 °C. Thereafter, the frozen cells were resuspended in ice-cold PBS supplemented with 10 mM imidazole and an EDTA-free protease inhibitor cocktail (Roche). Supernatants containing the nanobodies were collected by centrifugation at 20,000 g for 45 minutes at 4 °C. The His-tagged nanobodies were captured on an Ni^2+^-NTA affinity resin (Thermo Fisher Scientific) by gravity flow. Unbound proteins were washed away, and the nanobodies were eluted (PBS with 200 mM NaCl and 250 mM imidazole), after which the imidazole was removed by dialysis. The nanobody concentration was determined by measuring the absorbance at 280 nm in a NanoDrop (Thermo Fisher Scientific). From each stock solution 10 ug of nanobody was transferred, the buffer was exchanged, and the volume was reduced with SP3 bead cleanup [86] and following on-bead digestion. In brief, pure EtOH was added to a final concentration of 80%. 50 ug of each hydrophobic and hydrophilic beads (Cytiva, Sera-Mag™ Carboxylate-Modified [E7] Magnetic Particles Cat. #24152105050250 and Sera-Mag™ SpeedBead Carboxylate-Modified [E3] Magnetic Particles Cat. #65152105050250) were added to the solution, and incubated in a thermomixer at room temperature, 800 rpm, 15 minutes to allow binding. Samples were placed in a magnetic rack, and the solvent was removed. The remaining beads and bound proteins were washed 3 times with 90% EtOH, and were finally resuspended in 20 µL of 2.5 M guanidine hydrochloride (GuHCl; G3272 Sigma-Aldrich) and 250 mM HEPES solution (4-(2-Hydroxyethyl)piperazine-1-ethanesulfonic acid; 7365-45-9 Sigma-Aldrich). Nanobodies were reduced and alkylated with 10 mM TCEP (Tris(3-hydroxypropyl triazolyl methyl)amine; 762342 Sigma-Aldrich) and 40 mM CAA (2-Chloroacetamide; 79-07-2 Sigma-Aldrich), incubated for 10 minutes at 95 °C. Samples were diluted 5 times in MilliQ water, and 200 ng of trypsin (V5280 Promega Gold) was added to a 1:50 protease:proteome ratio, assuming no losses. Samples were digested overnight, at 37 °C, 450 rpm. The next day, samples were placed on a magnetic rack and the solution was transferred to a new tube. Approximately 500 ng of peptides, assuming no losses, were acidified and loaded on EvoTips with the standard loading protocol [87] for mass spectrometry analysis. The samples were analysed using the EvoSep One liquid chromatography platform, in line with an Orbitrap Exploris 480 mass spectrometer equipped with a FAIMSpro device.

Peptides were separated with a PepSep C18 column (15 cm x 75 µm, 1.9 µm PepSep, 1893473), over 31 minutes employing the Whisper100 40SPD method. Peptides were ionised with nanospray ionisation with an 10 µm emitter (PepSep, 1893527), and spray voltage of 2,300 V in positive ion mode, and ion transfer tube of 240 ℃. The total carrier gas flow was set to 3.6 L/min, and FAIMS was operated at standard acquisition. Spectra were acquired in data dependent resolution mode, under two different compensation voltages of -50 and -70, with identical settings. Cycle time was set to 2 seconds, with MS1 spectra acquired with 60,000 resolution, scan range of 375-1500, normalised AGC target of 300%, RF lens of 40%, and automatic injection time. Filters were set for peptide MIPS mode, inclusion of charge states 2-6, dynamic exclusion of 60 seconds with 10 ppm tolerance, and intensity threshold of 10,000. MS2 spectra were acquired with an isolation window of 1.6 m/z, normalised HCD of 30%, Orbitrap resolution of 30,000, first mass at 120 m/z, normalised AGC target of 100%, and automatic injection time. Data analysis was performed in Proteome Discoverer [88] v2.4, with Sequest HT [89] as the search engine. The database used was the *E. coli* reference proteome (Uniprot reviewed, UP000284592, 4,360 sequences, accessed 01/12/2022) concatenated with the nanobody sequences, and additional dynamic modifications of acetylation or methionine loss at the protein N-terminus, along with methionine oxidation, and static modification of carbamidomethylation. FDR control was performed with Percolator, at 1% and 5% target FDRs. Precursor quantification was performed with the Minora Feature Detector and Feature Mapper nodes in the processing and consensus workflows, respectively. Abundances were based on unique and razor peptides and above signal to noise ratio (S/N) > 5, and normalised based on total protein amount. PSMs at 1% FDR were exported for further processing, data extraction and model validation.

#### 4.4.2 HeLa proteome

HeLa cells were cultured in T25 flasks with Dulbecco’s Modified Eagle Medium (DMEM, Cat no 10565018, ThermoFisher Scientific) until confluency. Cells were pelleted with centrifugation, and resuspended in 6 M GuHCl. Proteins were reduced, alkylated and digested as for nanobodies above, with an additional LysC digestion for 1 h at 1:100 protease:protein ratio, before tryptic digestion. 200 ng of peptides, assuming no losses, were acidified and analysed with a nLC E1200 in line with an Orbitrap Exploris 480 mass spectrometer equipped with a FAIMSpro device. Peptides were separated with an 15 cm x 75 µm, 2 µm EASY-SpayTM column (ThermoFisher Scientific, ES904) over a 70 minute gradient, starting at 6% buffer B (80% ACN, 0.1% FA), increasing to 23% for 43 min, then to 38% for 12 min, 60% for 5 min, 95% for 3 min, and staying at 95% for 7 min. Peptides were ionised with electrospray ionisation with a positive ion spray voltage of 2,000, and ion transfer tube of 275 ℃. The rest of the method settings were as described above, with the difference of top 20 data dependent scans, and normalised HCD of 28% for MS2 spectrum acquisition. Data analysis was performed as above, with only differences being the use of human database (Uniprot reviewed, UP000005640, 20,518 sequences, accessed 05/03/2023), and lack of normalisation of precursor quantification in the consensus workflow.

#### 4.4.3 *S. brodae* proteome

Cells were pelleted and lysed under native conditions with hypotonic buffer (10 mM HEPES, 10 mM NaCl, 1.5 mM MgCl_2_, 2 mM EDTA, 0.1% NP-40, Roche Mini protease inhibitor) and a probe sonicator (20% power, 10 seconds with 1 second pulse, 5 rounds) on ice. Lysates were upconcentrated and buffer exchanged with spin filters (Amicon, 3 kDa cutoff, UFC500324, Merck Millipore) to 50 mM HEPES pH 7.8, and their concentration was determined by Nanodrop. From then on the standard proteomics sample preparation was followed, starting with 50 ug of proteome. Proteins were reduced, alkylated and digested as described above. Assuming no losses, 1 µg of peptides were acidified and loaded on EvoTips with the low input protocol. The samples were analysed with EvoSep One liquid chromatography platform, in line with an Orbitrap Eclipse mass spectrometer equipped with a FAIMSpro device. Peptides were separated with a PepSep C18 15 cm x 150 µm, 1.9 µm (PepSep, 1893471), over 44 minutes with the standard 30SPD method. Peptides were ionised with nanospray ionisation with an 10 µm emitter (PepSep, 1893527), and spray voltage of 2,300 V in positive ion mode, and ion transfer tube of 240 ℃. Spectra were acquired in data dependent acquisition mode, under two different compensation voltages of -50 and -70, with identical settings. Cycle time was set to 1.2 seconds, with MS1 spectra acquired with 60,000 resolution, and maximum injection time of 118 sec. MS2 spectra were acquired with an isolation window of 1.6 m/z, normalised HCD of 30%, with otherwise similar settings as above. Data analysis was performed as above, with only differences being the use of the putative proteome “*Candidatus Scalindua Brodae*” database, assembled from metagenomics data (Uniprot Trembl, UP000030652, 4,014 sequences, accessed 28/02/2023), and lack of normalisation of precursor quantification in the consensus workflow. In a secondary search, the raw data were searched against the *S. brodae* proteome as above, along with the proteomes of *K. stuttgartiensis* (UP000221734, 3,801 sequences, accessed 27/07/2023), *S. rubra* (UP000094056, 5,207 sequences, accessed 27/07/2023), and the *S. profunda* metagenome from a previous study (23,834 sequences) [90].

#### 4.4.4 GluC degradome and PRM monitoring

HeLa cell lysates were extracted as the HeLa proteome section. Six aliquots of 20 µg of lysate were resuspended in 100 mM HEPES, pH 7.8 to reduce the GuHCl concentration to 0.5 M. 200 ng of GluC endopeptidase (V1651, Promega) was added to three out of the six samples to a 1:100 ratio of protease:proteome, and all samples were incubated at 37 ℃, 450 rpm for 20 minutes. Samples were reduced, alkylated and digested with trypsin as described previously. The next day, volume equivalent to 1 µg from each sample, assuming no losses, was loaded on EvoTips as described above, and samples were analysed using the EvoSep One liquid chromatography platform, in line with an Orbitrap Eclipse mass spectrometer equipped with a FAIMSpro device. Peptides were eluted from a PepSep C18 column (15 cm x 75 µm, 1.9 µm PepSep, 1893473) over 58 minutes with the Whisper100 20SPD method. Scans were acquired with the same settings as in the HeLa proteome single shot analysis. Data analysis was performed as above, with use of the human database for the HeLa proteome searches, semi-tryptic search, and precursor quantification normalised on the total peptide amount from each sample in the consensus workflow.

PRM assays were designed for representative peptides detected by Instanovo with high confidence, but not with the database search. Peptide sequences were imported in Skyline [91], and an inclusion list with the precursor masses was exported. The inclusion list was used to create a PRM monitoring method with a targeted mass inclusion filter for acquisition of MS/MS scans. GluC degradome samples were analysed with the same setup as in shotgun proteomics and the same FAIMS CVs. Scans were acquired with 60,000 resolution for MS1, and 15,000 resolution for MS2 and a cycle time of 1 sec for each FAIMS CV, with otherwise similar settings with the shotgun proteomics experiment. Results were analysed and visualised with Skyline.

#### 4.4.5 External dataset analysis

The raw data from snake venom proteomics dataset were downloaded and reanalyzed using the Uniprot database sequences for the serpentes order (331,759 sequences, accessed 05/09/2022), similar to the original study. Data were analysed with Proteome Discoverer v2.4 and the Sequest HT search engine, with all files included in the same analysis, normalisation on total peptide amount and precursor quantification, with other settings similar to other datasets. The herceptin dataset was downloaded and analysed similarly. However, the raw data from the 6 different proteases were searched separately, and no precursor or normalisation was performed. The same fasta database as in the original study was used for PSM detection. Search results were then combined for prediction and evaluation.

The immunopeptidomics dataset was reprocessed with the same proteome database as in the original paper with MSFragger[92] and the FragPipe v21.1 pipeline with the non-specific HLA workflow, and otherwise default settings. MSBooster[93] was used for rescoring with deep learning prediction, and Percolator was used for PSM FDR control, while no FDR control was used on the protein level.

The wound fluid dataset was downloaded and searched with the same human database as used for the HeLa proteome and GluC degradomics experiments. Both raw data files were analysed in the same search in Proteome Discoverer v2.4, with total peptide amount normalisation and precursor quantification. In the secondary search results, the same human proteome as well as protein sequences downloaded from the Uniprot database for the pathogens of interest *Citrobacter sp.* (UP000682339, 3,414 sequences), *P. aeruginosa* (UP000002438, 5,564 sequences), *S. aureus* (UP000008816, 2,889 sequences), and *E. coli* (UP000000625, 4,403 sequences) were used for PSM detection.

#### 4.4.6 Analysis rationale

During dataset preprocessing, we filter out spectra with more than 800 peaks. In our labelled datasets, we remove PSMs with more than 30 residues length. Both of these constraints were set due to the necessity for determining a maximum number of peaks and maximum number of residues in peptide sequences for the input and output vector sizes of our models respectively. These cutoffs were selected based on observed distributions of these attributes in the ACPT dataset from Proteome Tools, although the dataset consisted of pools of synthetic peptides. We are aware that by setting these thresholds, we are missing a significant number of spectra and PSMs from our biological datasets, as well as any potential testing datasets to come. Especially in data acquisition runs where the spectra have higher noise, i.e. contains higher interference and contaminants, or in cases where MS2 spectra are acquired in profile mode, the strict cutoff of 800 peaks might be too low. To alleviate these issues in future studies, we are planning on selecting the top 800 peaks from each MS2 spectrum, instead of completely removing them from the dataset (as we did in this study for the full search space datasets). A prime example of this noisiness was observed in the immunopeptidomics dataset reanalysed in this study, where only 651 spectra passed this peak threshold. Therefore, only the full database search results and the prediction for all scans in that dataset was used for downstream analysis. In the future, a model with double the number of maximum output residues (60 instead of 30) is envisioned, which will cover the vast majority of bottom up applications, and a larger model can be considered for top down applications if deemed possible.

When reporting protein numbers, unless otherwise specified, we talk about protein groups, meaning the number of candidate groups that peptides map to, or the set of the first protein identifiers for each peptide mapping. Protein inference, grouping, and master protein assignment was out of the scope of this study.

We regard the residues isoleucine and leucine as interchangeable when inferring and mapping predicted sequences to proteins, as these amino acids are isobaric. Although it might be possible through the detection of w ions for the model to distinguish between the two isobaric amino acids, this differentiation is not yet possible with our model and fine tuning or improving on this will be part of future efforts. Protein coverage was calculated as the number of amino acids covered by PSMs or predictions, by non-redundant peptide sequences, expressed as a percentage.

To evaluate FDR in our *de novo* peptide sequencing predictions, we ground our model’s confidence with the FDR estimation of the associated database searches. This means that we extract the scans that we have a PSM in the database search, and compare these with our predictions for the same scans, ranked based on model confidence. Once we have reached a 5% FDR threshold, as decided by the PSM-prediction comparison (in this case, the database search is our ground truth, and the PSMs in it our correct labels), we regard this confidence threshold as our model’s threshold for 5% FDR. We are aware of the limitations of this approach, and are working on better assessment criteria for FDR in *de novo* peptide sequencing.

## Data availability

Raw data and search results used for evaluation, for public datasets used or datasets generated in this study, have been deposited to the ProteomeXchange Consortium via the PRIDE [94] partner repository with the dataset identifier PXD044934. The repository can be accessed with reviewer account username and password Tf8dE6nv. Additional files relating to pre-processed results used for training and metric evaluation have also been uploaded in the same archive repository. Supplementary files supporting the data pre-processing, tool usage, and analysis performed on 8 different application-centric datasets have been deposited on figshare (10.6084/m9.figshare.24173889.

The ProteomeTools datasets used to train the models in this study can be found in the PRIDE repository with identifiers PXD004732 (Part I), PXD010595 (Part II), and PXD021013 (Part III). The nine species dataset [33] is available through the MassIVE repository with dataset identifier MSV000081382. The immunopeptidomics dataset used for model evaluation can be found in the PRIDE repository with identifier PXD006939. Snake venom files and search results can be found in the PRIDE repository with identifier PXD036161. The wound exudate files and search results are available in PanoramaWeb with dataset identifier PXD025748. The herceptin dataset is available in Figshare (doi.org/10.6084/m9.figshare.21394143).

## Code availability

InstaNovo is available at github.com/instadeepai/InstaNovo along with model checkpoints. Here, we have also implemented a Google Colab notebook test version for easy access. Furthermore, we have made the nine-species dataset [33] also available at huggingface.co/datasets/InstaDeepAI/ms_ninespecies_benchmark, and the high-confidence ProteomeTools [39] dataset available at huggingface.co/datasets/InstaDeepAI/ms_proteometools. Custom scripts used for data analysis and most of visualisation are available upon request. A hosted API application for analysis of high throughput, large scale datasets was outside the scope of this study, but is planned for the future.

## Footnotes

1 In all of our experiments, we used residues with the following post-translational modifications: carbamidomethylation for cysteine, oxidation for methionine, and deamidation for asparagine and glutamine.

2 Our system also returns the sequence of per-residue log-probabilities log p(y_*i*_|y_*<i*_, **m, I**, m_*prec*_, c_*prec*_) for quantifying uncertainty around predicted residues in downstream analysis.

## Acknowledgements

We extend our gratitude to the DTU Proteomics Core facility staff for calibration and maintenance of instruments. We especially thank Maike Wennekers Nielsen and Marie Vestegaard Lukassen for their help and advice during troubleshooting, sample preparation and data acquisition. *S. brodae* reactor co-enrichment cultures were a kind gift of Laura van Niftrik, Sebastian Lücker, and Guylaine Nuijten from the Department of Microbiology, Faculty of Science, Radboud University, Nijmegen (Netherlands). Models were trained on InstaDeep’s AIchor computing cluster (aichor.ai). Biorender (biorender.com) was used to generate parts of the graphical abstract and the figures in this manuscript.

This work is dedicated to the memory of Prof. Dr. Ulrich auf dem Keller. His knowledge, insights, and directions were instrumental to our research, and he is deeply missed.

## Author information

Authors and affiliations

Kevin Eloff

**InstaDeep, 5 Merchant Square, London, UK**

Konstantinos Kalogeropoulos

**Department of Biotechnology and Biomedicine, Technical University of Denmark, Kongens Lyngby, Denmark**

Oliver Morell

**Department of Biotechnology and Biomedicine, Technical University of Denmark, Kongens Lyngby, Denmark**

Amandla Mabona

**InstaDeep, 5 Merchant Square, London, UK**

Jakob Berg Jespersen

**Department of Biotechnology and Biomedicine, Technical University of Denmark, Kongens Lyngby, Denmark; Novo Nordisk Foundation Center for Biosustainability, Technical University of Denmark, Kongens Lyngby, Denmark**

Wesley Williams

**InstaDeep, 5 Merchant Square, London, UK**

Sam van Beljouw

**Department of Bionanoscience, Delft University of Technology, 2629 HZ Delft, Netherlands; Kavli Institute of Nanoscience, 2629 HZ Delft, Netherlands**

Marcin Skwark

**InstaDeep, 5 Merchant Square, London, UK**

Andreas Hougaard Laustsen

**Department of Biotechnology and Biomedicine, Technical University of Denmark, Kongens Lyngby, Denmark**

Stan J. J. Brouns

**Department of Bionanoscience, Delft University of Technology, 2629 HZ Delft, Netherlands; Kavli Institute of Nanoscience, 2629 HZ Delft, Netherlands**

Anne Ljungars

**Department of Biotechnology and Biomedicine, Technical University of Denmark, Kongens Lyngby, Denmark**

Erwin Schoof

**Department of Biotechnology and Biomedicine, Technical University of Denmark, Kongens Lyngby, Denmark**

Jeroen Van Goey

**InstaDeep, 5 Merchant Square, London, UK**

Ulrich auf dem Keller

**Department of Biotechnology and Biomedicine, Technical University of Denmark, Kongens Lyngby, Denmark**

Karim Beguir

**InstaDeep, 5 Merchant Square, London, UK**

Nicolas Lopez Carranza

**InstaDeep, 5 Merchant Square, London, UK**

Timothy Patrick Jenkins

**Department of Biotechnology and Biomedicine, Technical University of Denmark, Kongens Lyngby, Denmark**

## Contributions

KK and TPJ conceived the research idea. KE designed the model with input from KK, NC, JVG, WW, OM, MS, KB, and TPJ. KK, TPJ, AL, AHL, ES, and UK selected the validation datasets. KK performed sample preparation and mass spectrometry analysis. KK, JJ, and AL prepared these and additional public datasets and KK, KE, AM, and OM analysed them. KK, KE, TPJ, AM, WW, and OM wrote the manuscript with input from all authors. All authors read and approved the final version of the manuscript.

The two first authors contributed equally and the first name was selected randomly, as such both first authors have the permission of all co-authors to cite their name first in their CV’s and any other official correspondence.

## Corresponding authors

Correspondence and requests for materials should be addressed to T.P.J., K.K., and K.E.

## Ethics declarations

KE, AM, JVG, WW, MS, KB, and NC are employees of InstaDeep, 5 Merchant Square, London, UK. The remaining authors declare no conflicts of interest.

## Supplementary

**Supplementary Fig. 1.**
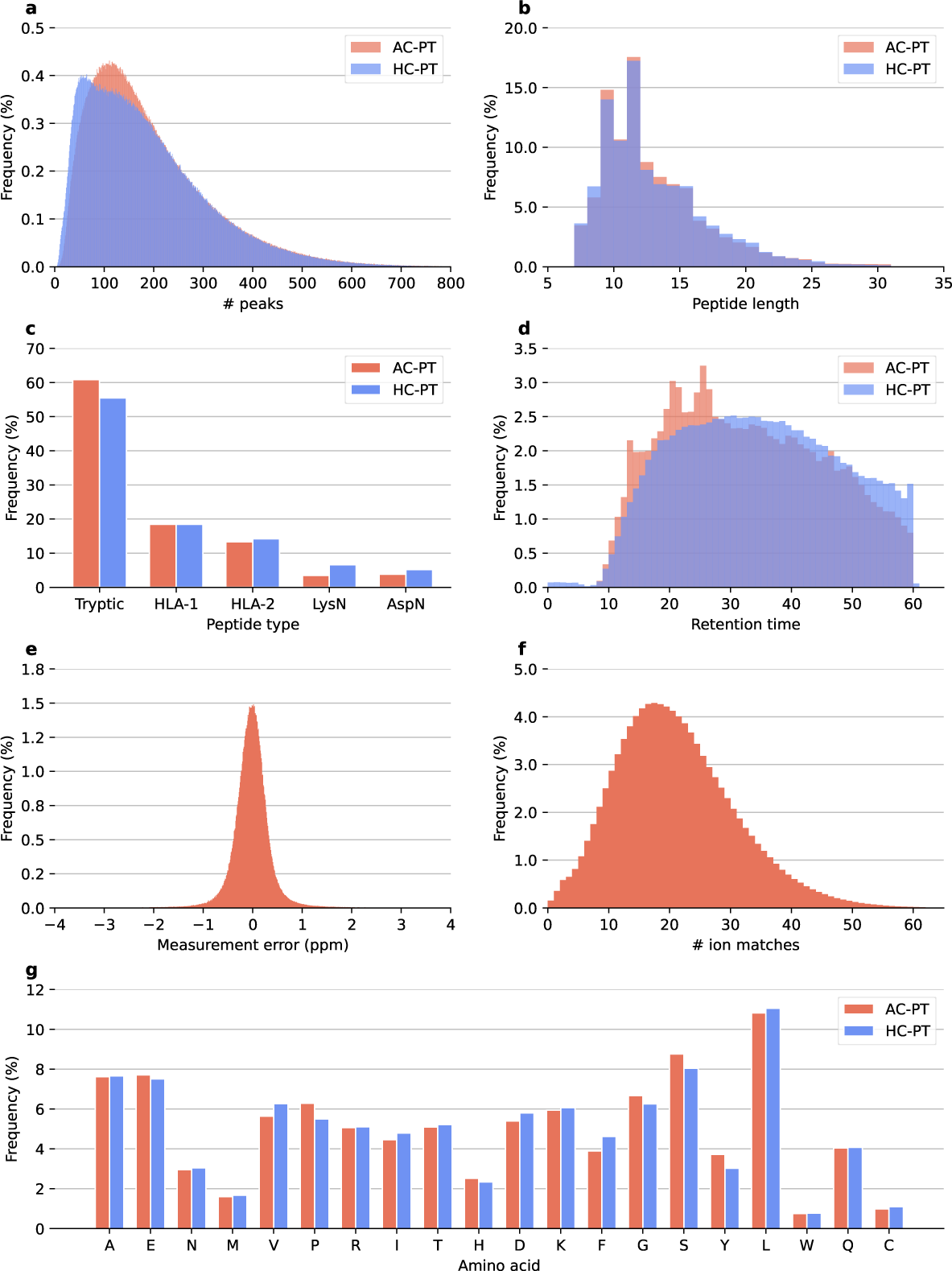
ProteomeTools descriptive statistics for all-confidence PSMs (AC-PT) and high-confidence PSMs (HC-PT). **a,** Number of peaks per spectra. **b,** Peptide length of PSM sequences. **c,** Distribution by peptide type, including tryptic, HLA-I, HLA-II, LysN, and AspN. **d,** Retention time distribution. **e,** Distribution of measurement error (ppm) in AC-PT. **f,** Ion matches for PSM scans distribution in AC-PT. **g,** Amino acid frequency in PSM sequences.

**Supplementary Fig. 2.**
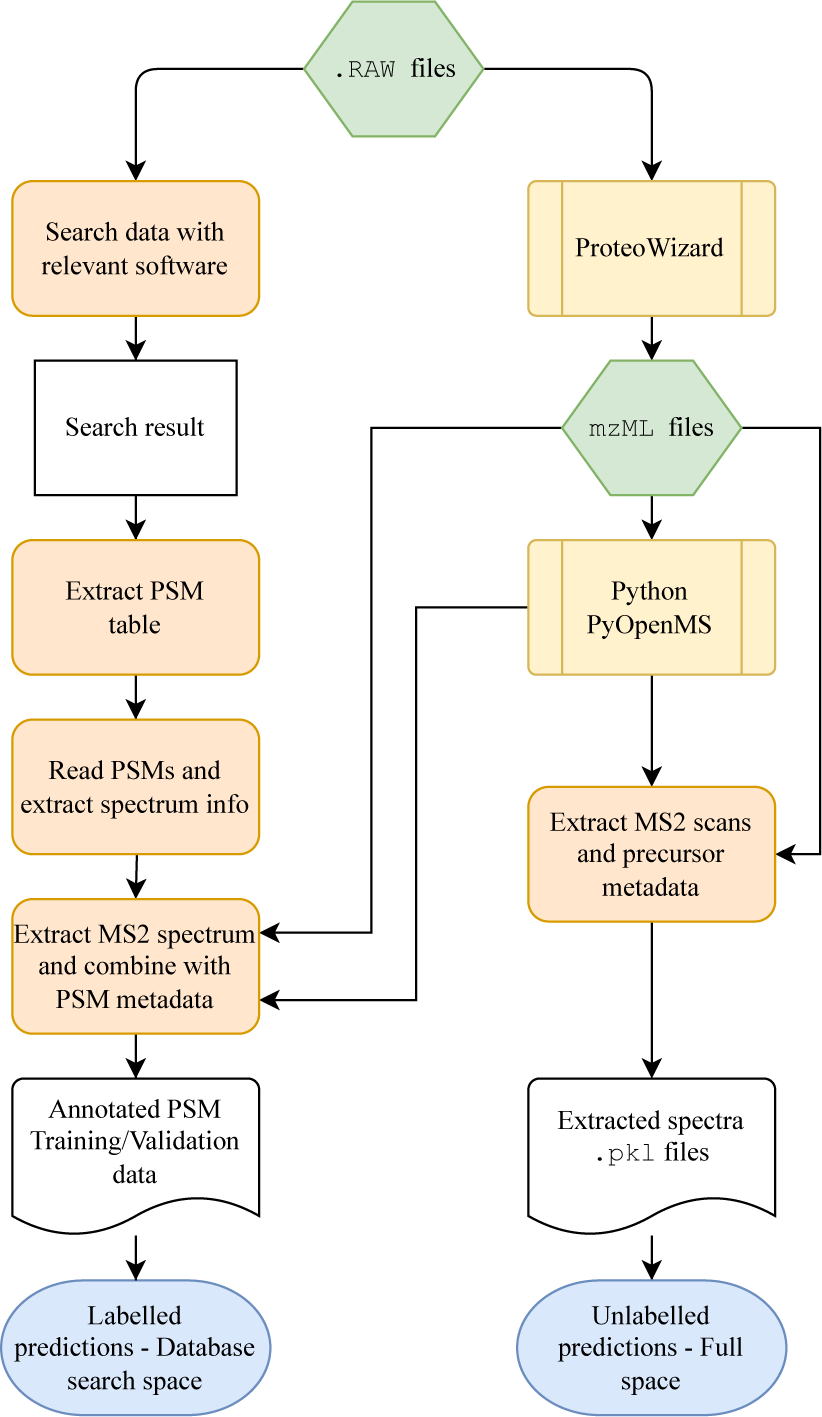
Detailed workflow for dataset extractions and preprocessing. Thermo .raw files were searched against a proteome database with Proteome Discoverer or MaxQuant. MSConvert from the Proteowizard tool suite was used to convert .raw files to mzML. The mzML files were used to extract m/z and intensity vectors as well as associated metadata with pyOpenMS. Two datasets were created, the database search space containing only scans that were matched to peptide hits (PSMs) from the database searches, and the full search space which contained all MS/MS scans from all .raw files of each experiment.

**Supplementary Fig. 3.**
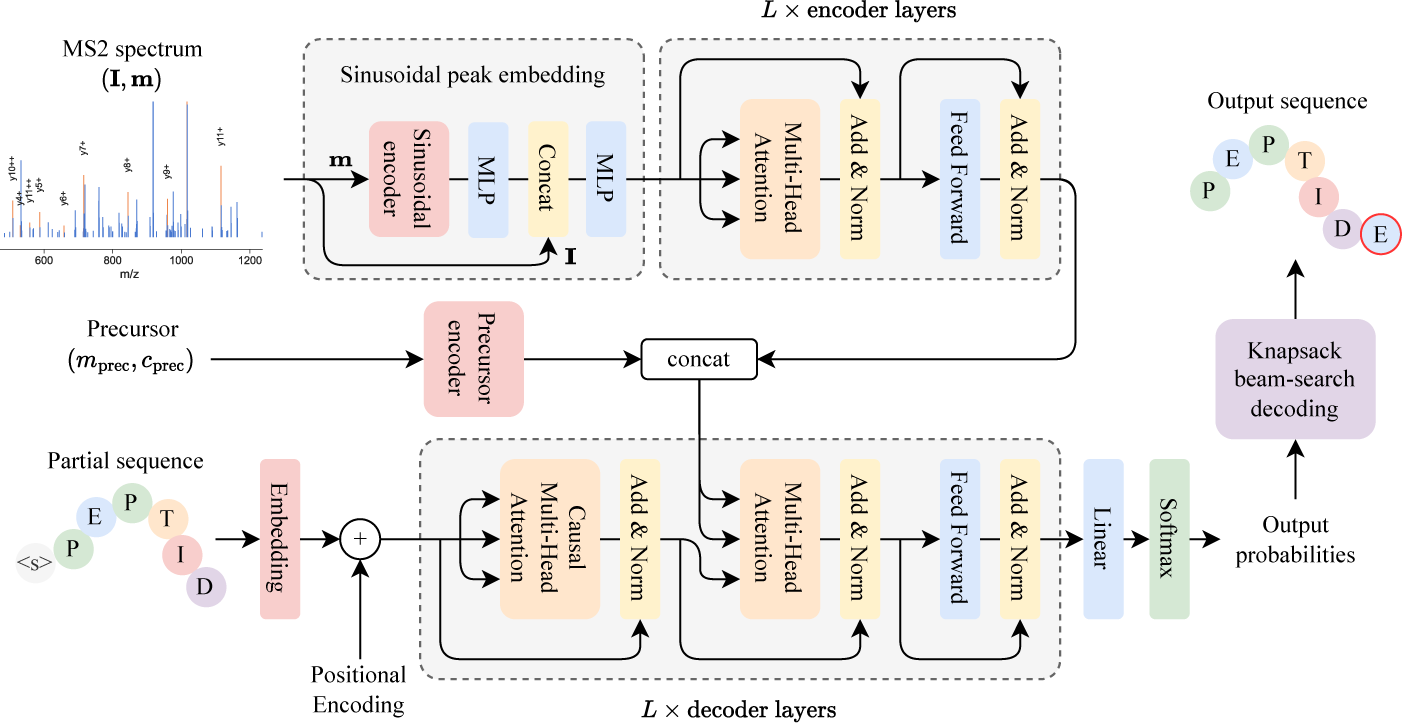
Detailed model architecture and description of InstaNovo model. The precursor information may also be provided as a start-of-sequence embedding at the input to the decoder layers.

**Supplementary Fig. 4.**
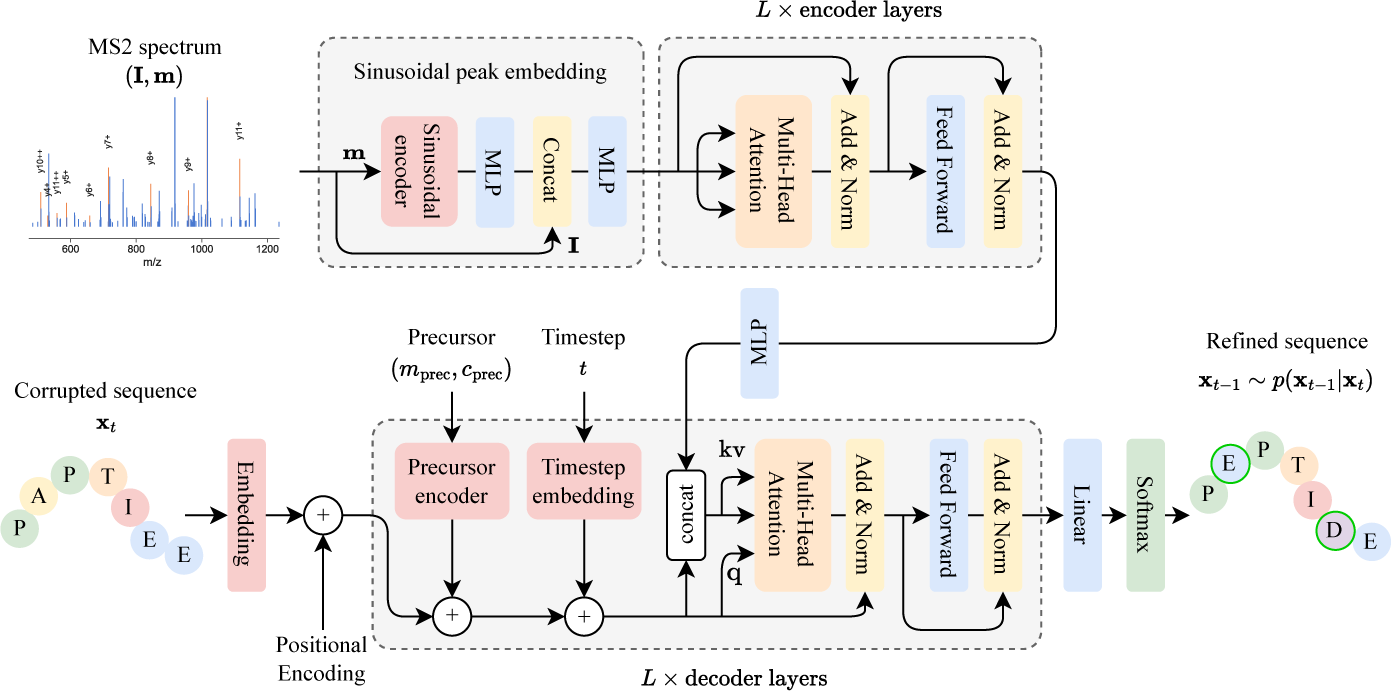
**Detailed model architecture and description of InstaNovo+ model.**

**Supplementary Fig. 5.**
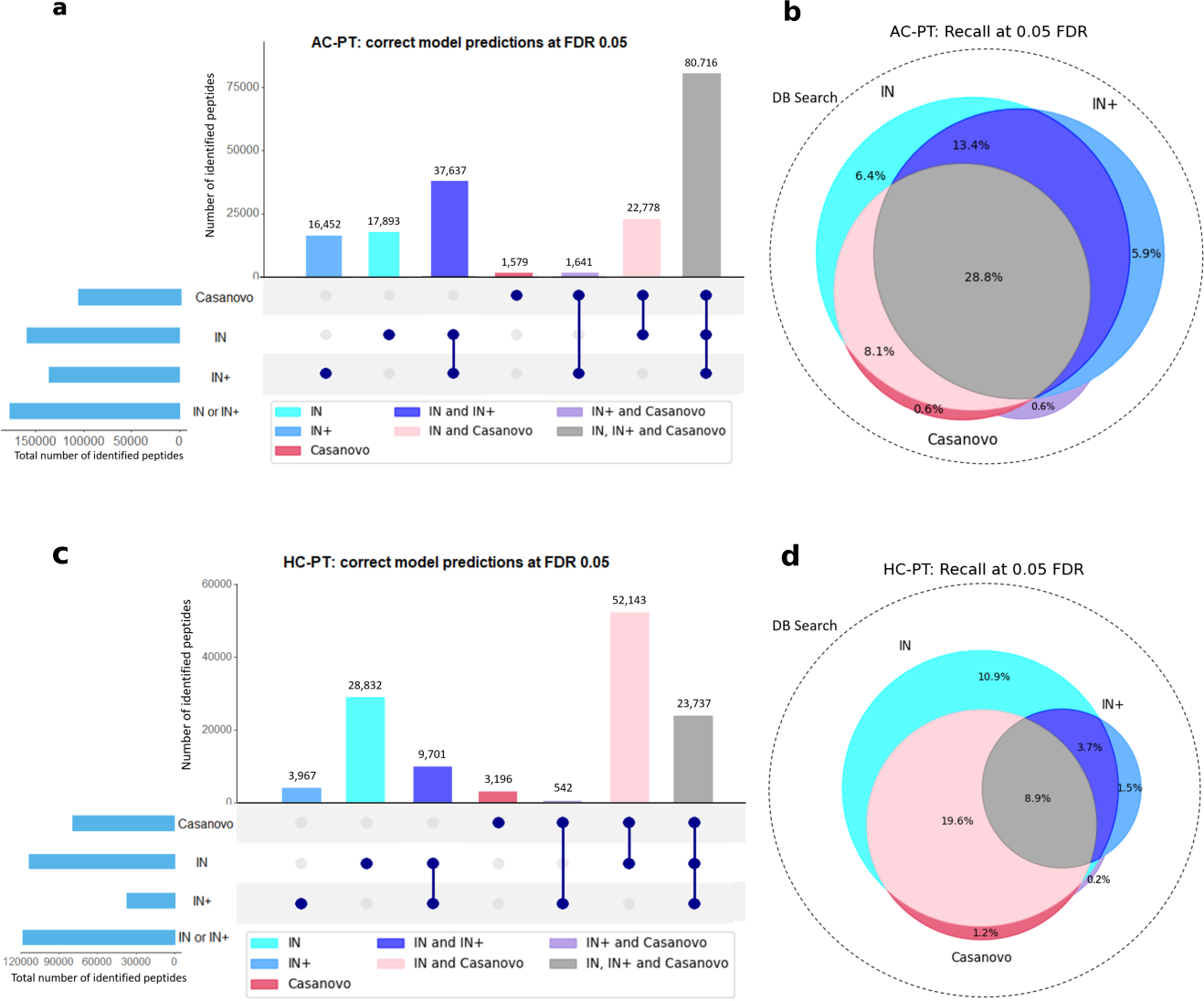
Overlaps between IN, IN+ and Casanovo’s correct predictions at 0.05 FDR for AC-PT and HC-PT. **a,** Peptide-level UpSet plot illustrating the intersection of correct predictions made by the IN, IN+, and Casanovo models on the AC-PT dataset, when evaluated at a false discovery rate (FDR) of 0.05. **b,** Peptide-level Venn Diagram illustrating the same intersections as figure a, but displaying them as percentages (recall) of the DB search ground truth dataset, which is illustrated by the area of the circle with the dotted edge. Areas in the Venn diagram are approximate, due to the imperfection of the Venn algorithm. **c,** Equivalent of figure a, for the HC-PT dataset **d,** Equivalent of figure b, for the HC-PT dataset

**Supplementary Fig. 6.**
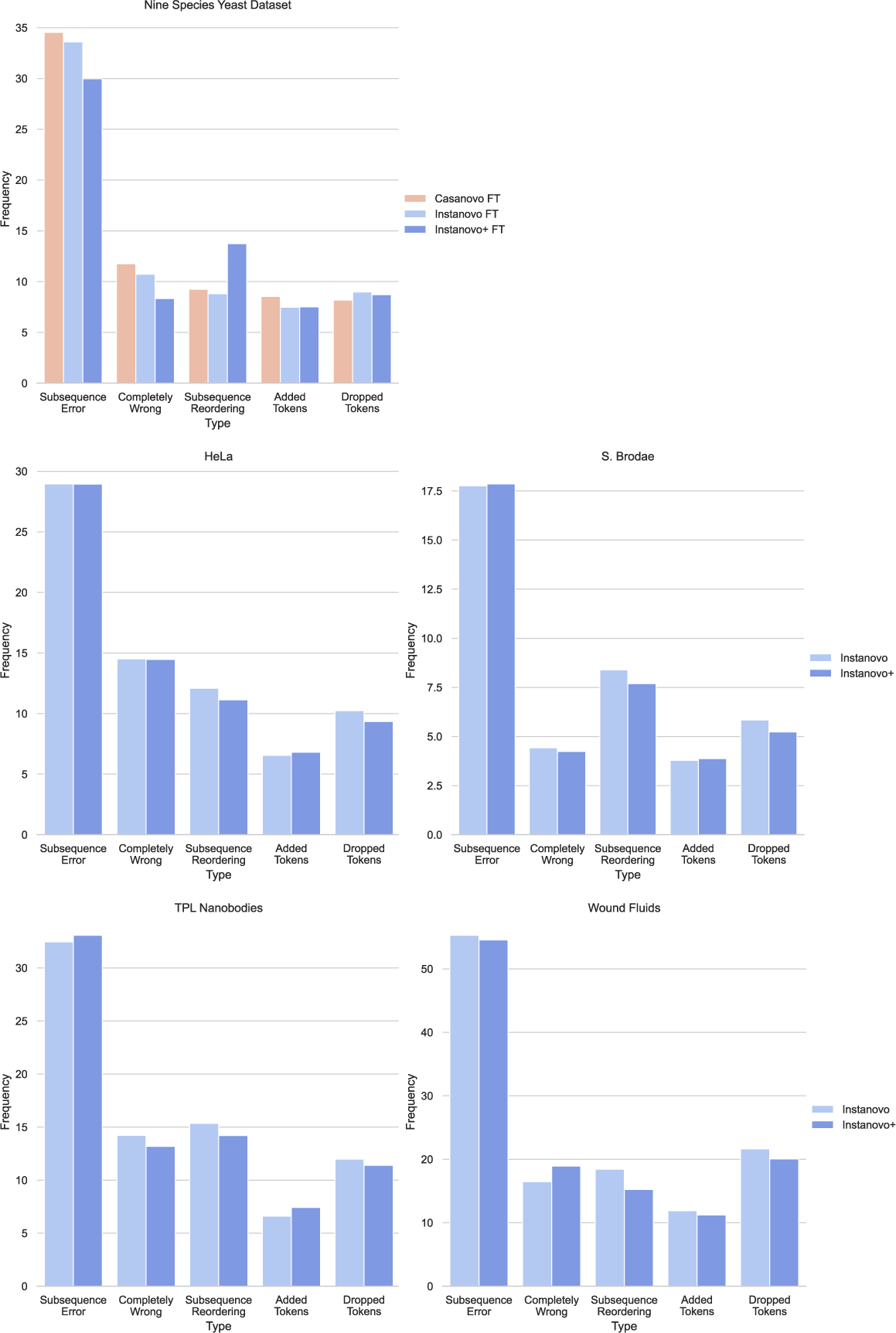
Error analysis for a selection of evaluation datasets. Top left: Comparison of Casanovo, IN, and IN+ predictions errors in the nine-species dataset. Most errors are caused by a few errors in the overall amino acid sequence for all models. Bottom: Comparison of IN and IN+ errors in 4 out of the 8 biological datasets.

**Supplementary Fig. 7.**
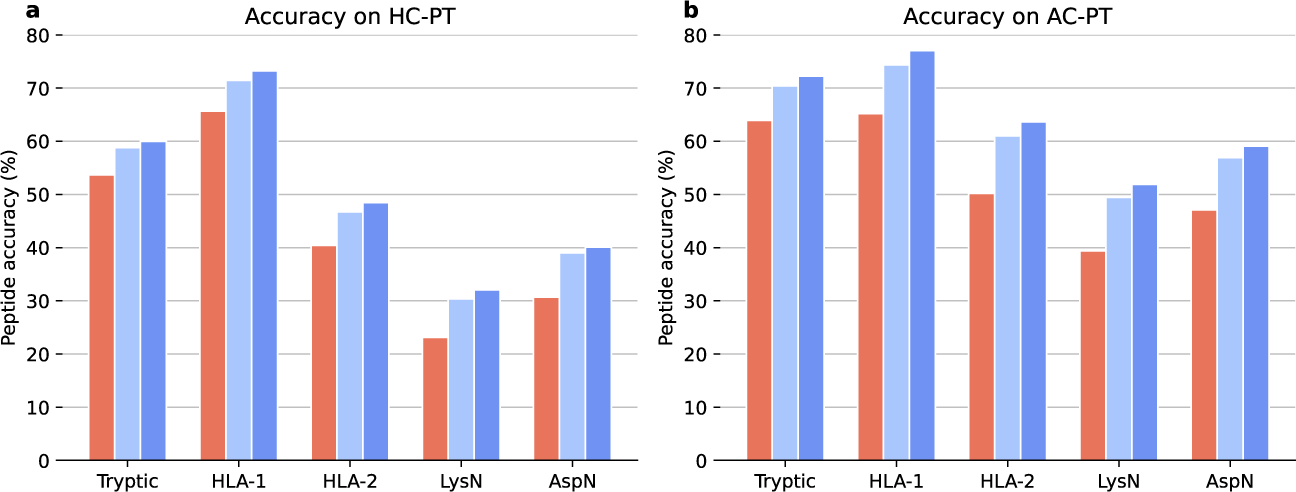
**Peptide accuracy on HC-PT and AC-PT grouped by the type of peptide.**

**Supplementary Fig. 8.**
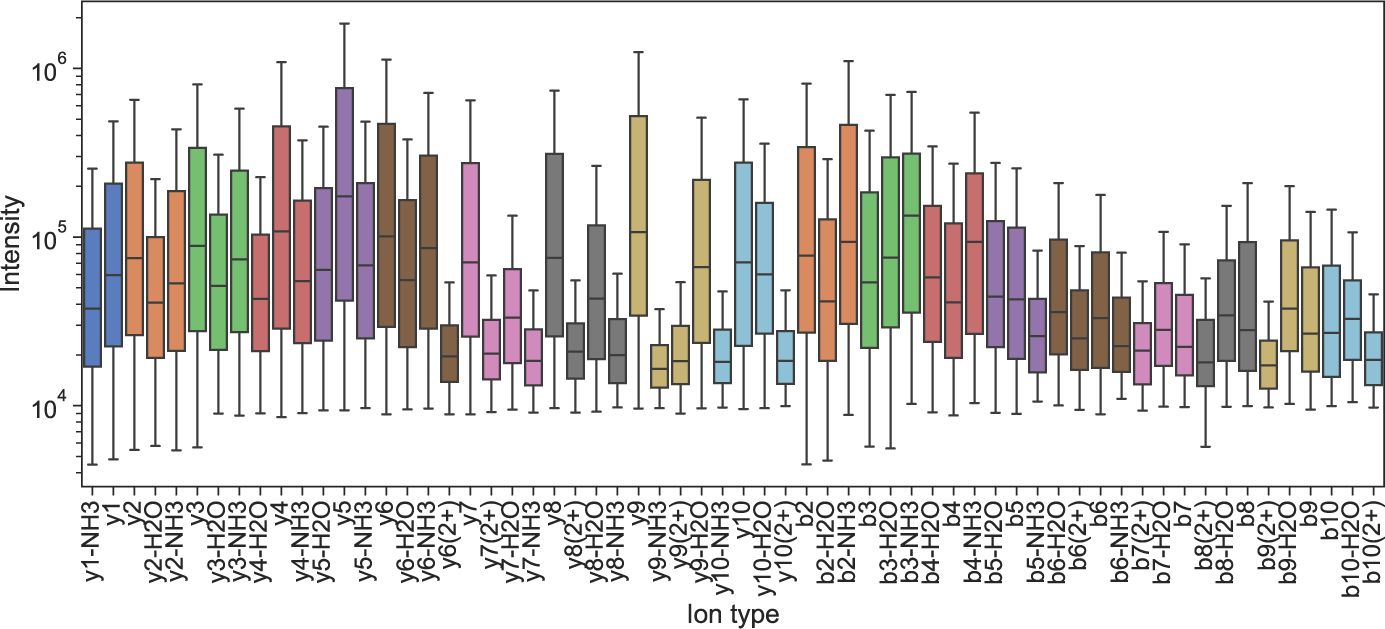
**Fragment ion log intensity from a selected analytical run from the training data, showing all ion types from identified PSMs.**

**Supplementary Fig. 9.**
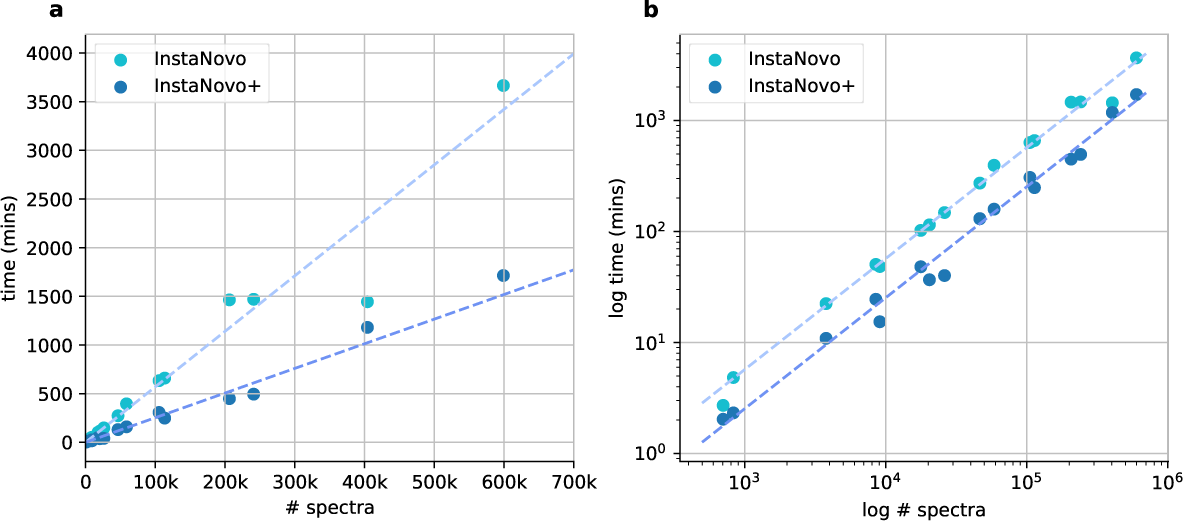
Runtime analysis of InstaNovo and InstaNovo+ showing a linear relationship. **a,** Inference time in minutes compared to the number of spectra. **b,** Comparison of runtime in the log-domain. The dashed lines in **a** and **b** depict the mean runtime per spectra. Presented runtimes were performed on hardware setup B (Supplementary Table 4). Hardware setup A saw a 2*×* average runtime improvement over setup B in knapsack decoding, due to poor performance of the server-grade CPU becoming a bottleneck in beam-search knapsack. When using InstaNovo predictions as a starting point for InstaNovo+, the total runtime would be the sum of the two models.

**Supplementary Fig. 10.**
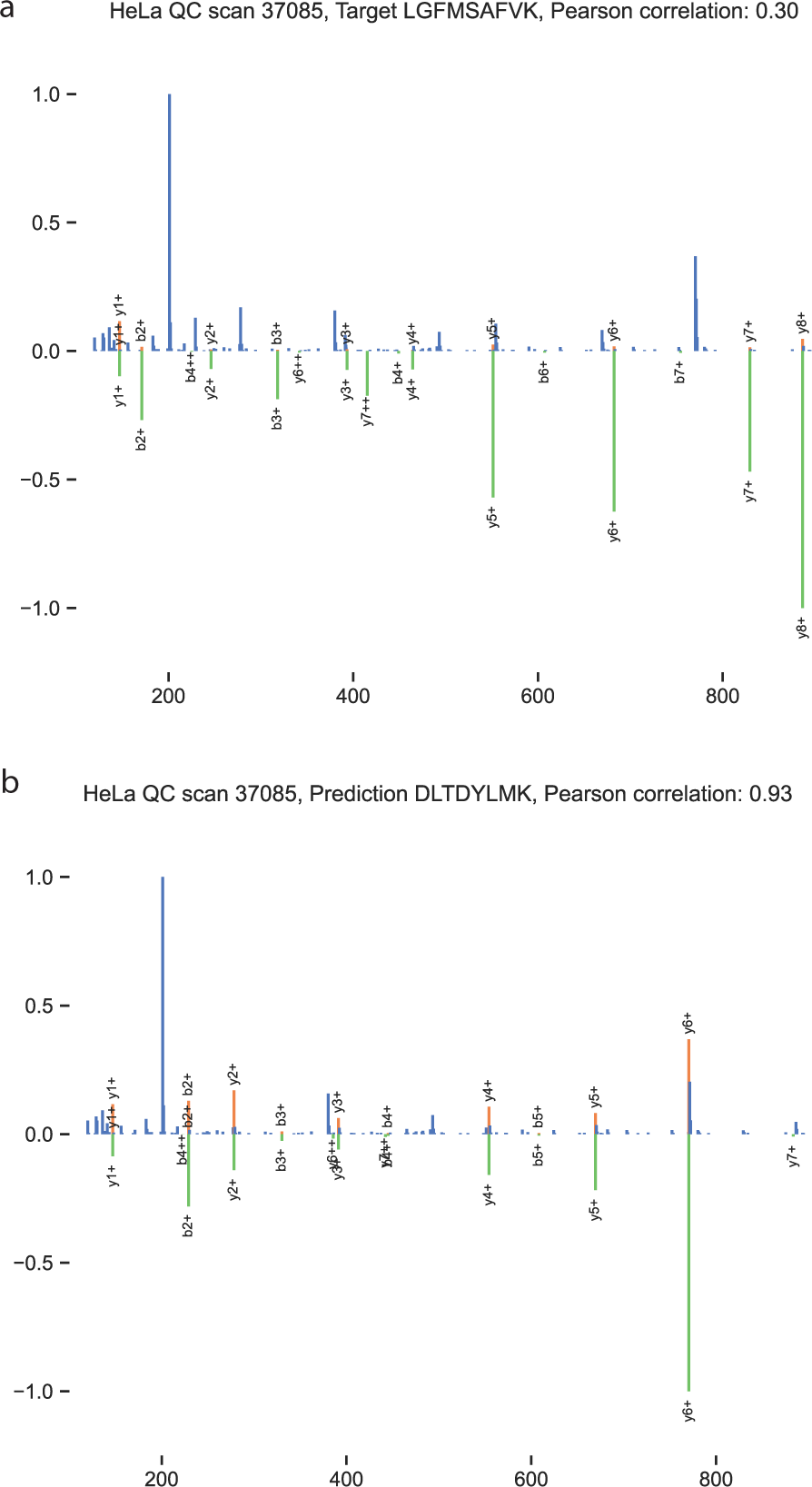
InstaNovo predicted sequence correlates better with observed spectrum than with database search PSM. **a,** Mirror plot for database search PSM sequence LGFMSAFVK in HeLa QC dataset, scan number 37,085. Top, experimental spectrum, bottom, Prosit predicted spectrum for the same sequence. **b,** Similar plot for predicted sequence, exhibiting higher similarity with experimental spectrum.

**Supplementary Fig. 11.**
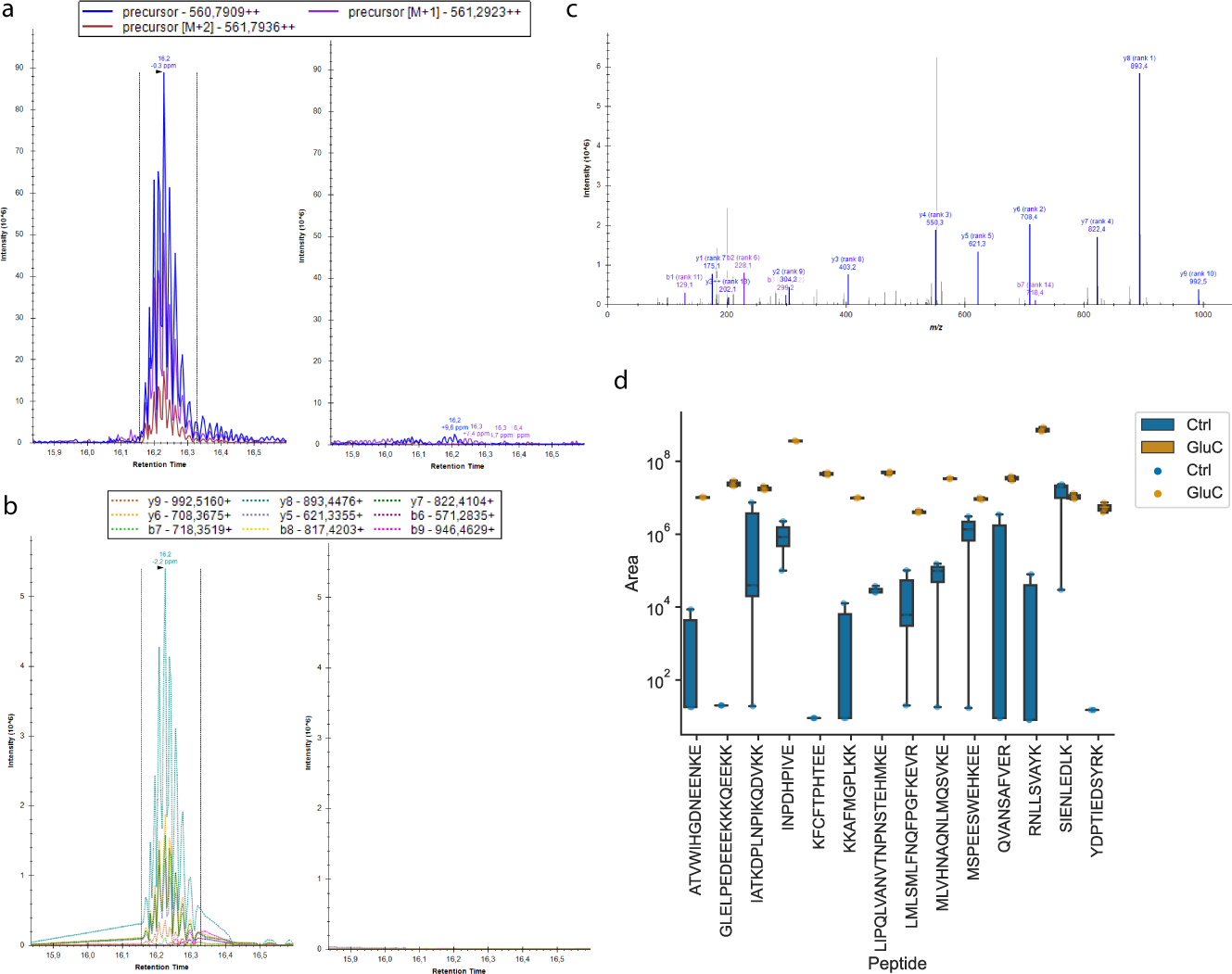
Targeted proteomics in GluC dataset. **a,** Monitoring of precursor mass for peptide QVANSAFVER in one GluC digested (left) and control (right) replicates. **b,** Monitoring of peptide transitions with fragment ion masses for the same peptide and replicates. **c,** Experimental spectrum at the apex of the transition peaks in the GluC digested replicate for the same peptide. **d,** Boxplot and striplot visualisation of the sum of fragment ion areas for selected peptides monitored with targeted proteomics.

**Supplementary Fig. 12.**
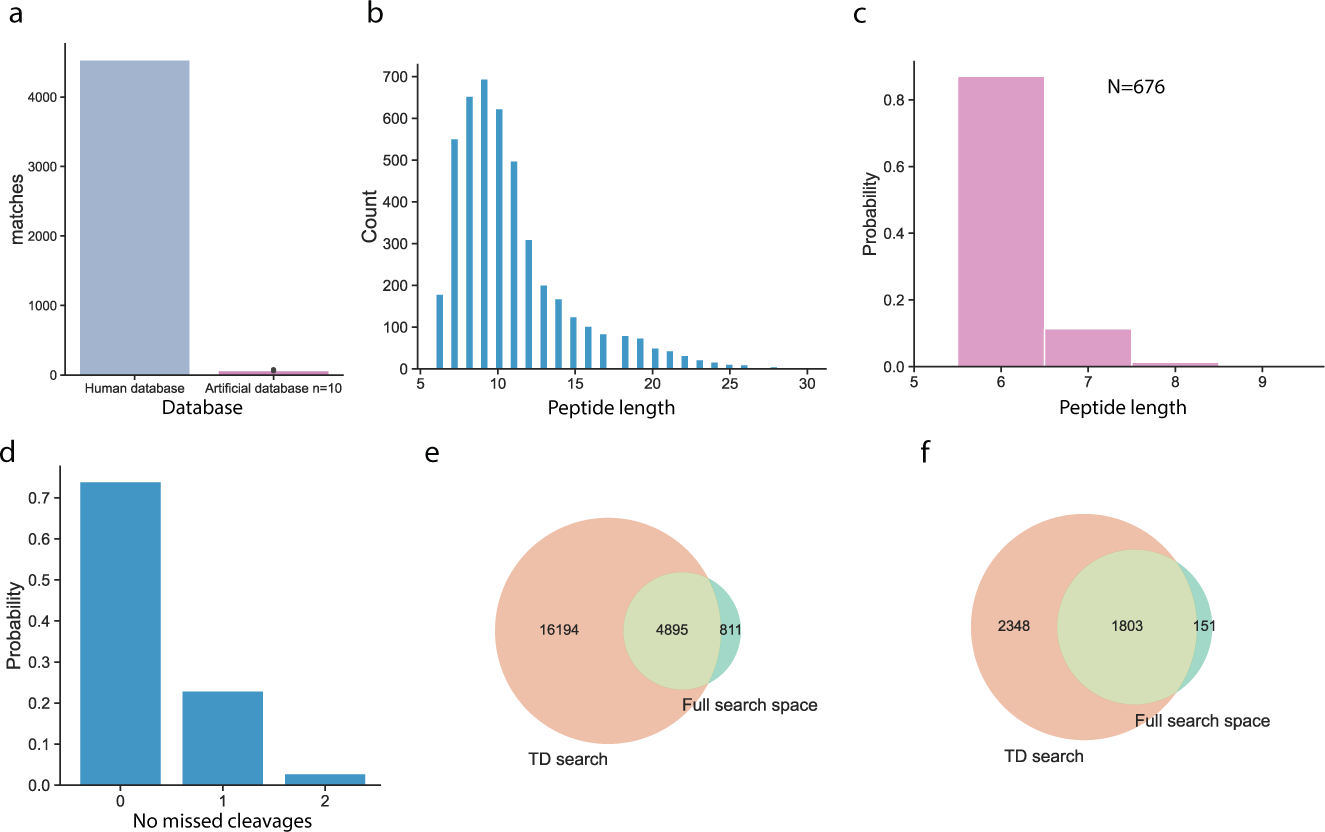
Supplementary figure for HeLa proteome analysis. **a,** Human database vs artificially generated database peptide matches comparison, database search space. **b,** Peptide length distribution in human proteome mapped predictions. **c,** Length of prediction matches in 10 artificially and randomly generated databases. **d,** Distribution of missed cleavages in full space predictions at 5% FDR. **e,** Venn diagram of peptide sequences mapping to the human proteome, identified with database search and sequences predicted by Instanovo in the full search space. **f,** Proteins identified from peptide sequences of database search PSMs or InstaNovo predictions in the full search space at 5% FDR.

**Supplementary Fig. 13.**
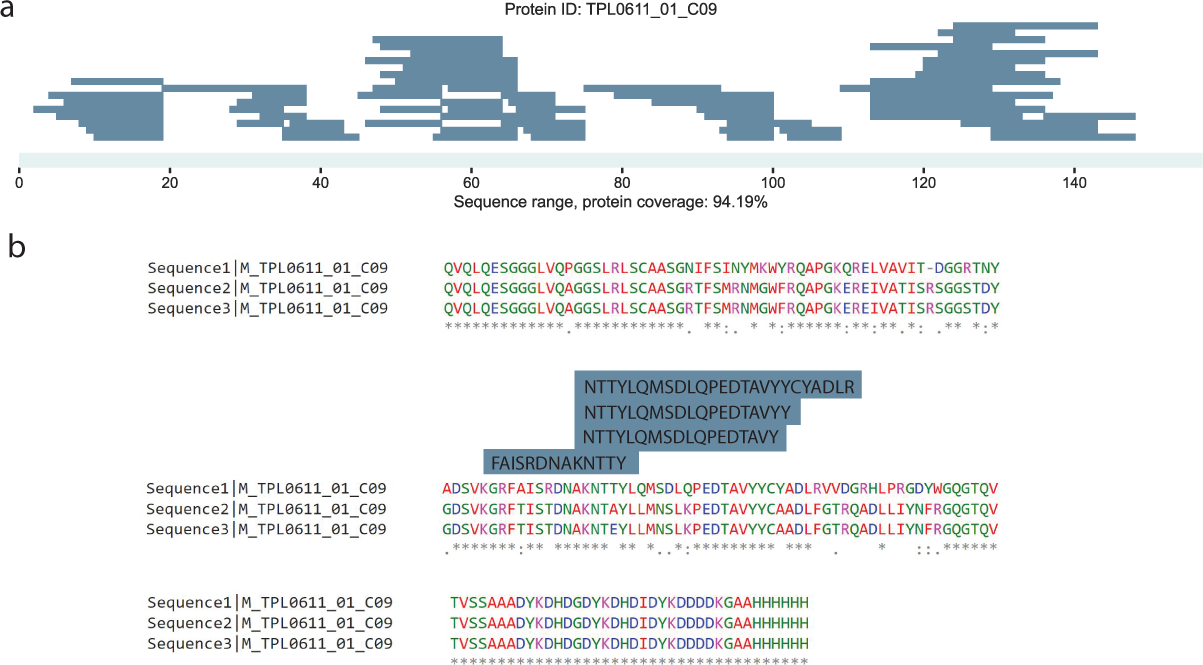
Direct sequencing and conflict resolution with InstaNovo. **a,** Nanobody TPL0611_01_C09 coverage and sequencing depth with unique peptides predicted at 5% FDR. **b,** Alignment of three separate sequencing runs on cells expressing the C09 nanobody, annotated with unique peptide sequences predicted with InstaNovo, mapping to one of the areas where there was ambiguity in determination of the sequence with genome sequencing methods.

**Supplementary Fig. 14.**
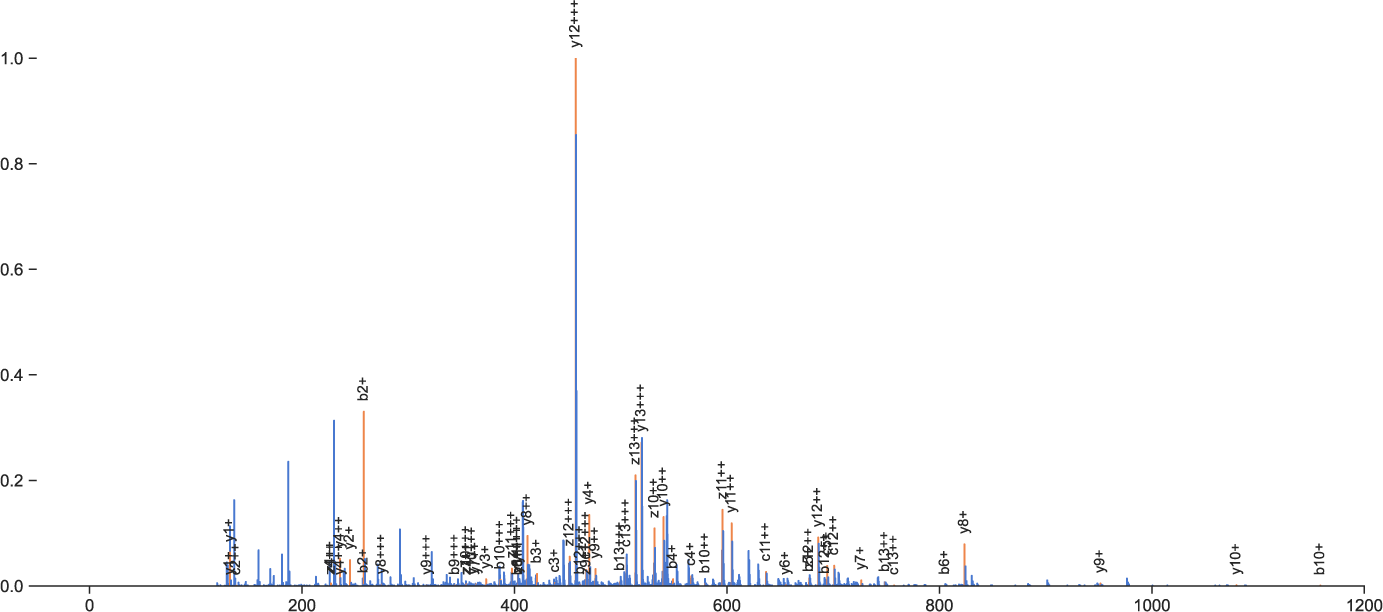
InstaNovo detects peptides from different fragmentation schemes. Spectrum of peptide sequence AWYQQKPGKAPKLL, assigned correctly by Instanovo in the Herceptin light chain. The spectrum was generated with EThcD fragmentation from the elastase digested antibody samples.

**Supplementary Fig. 15.**
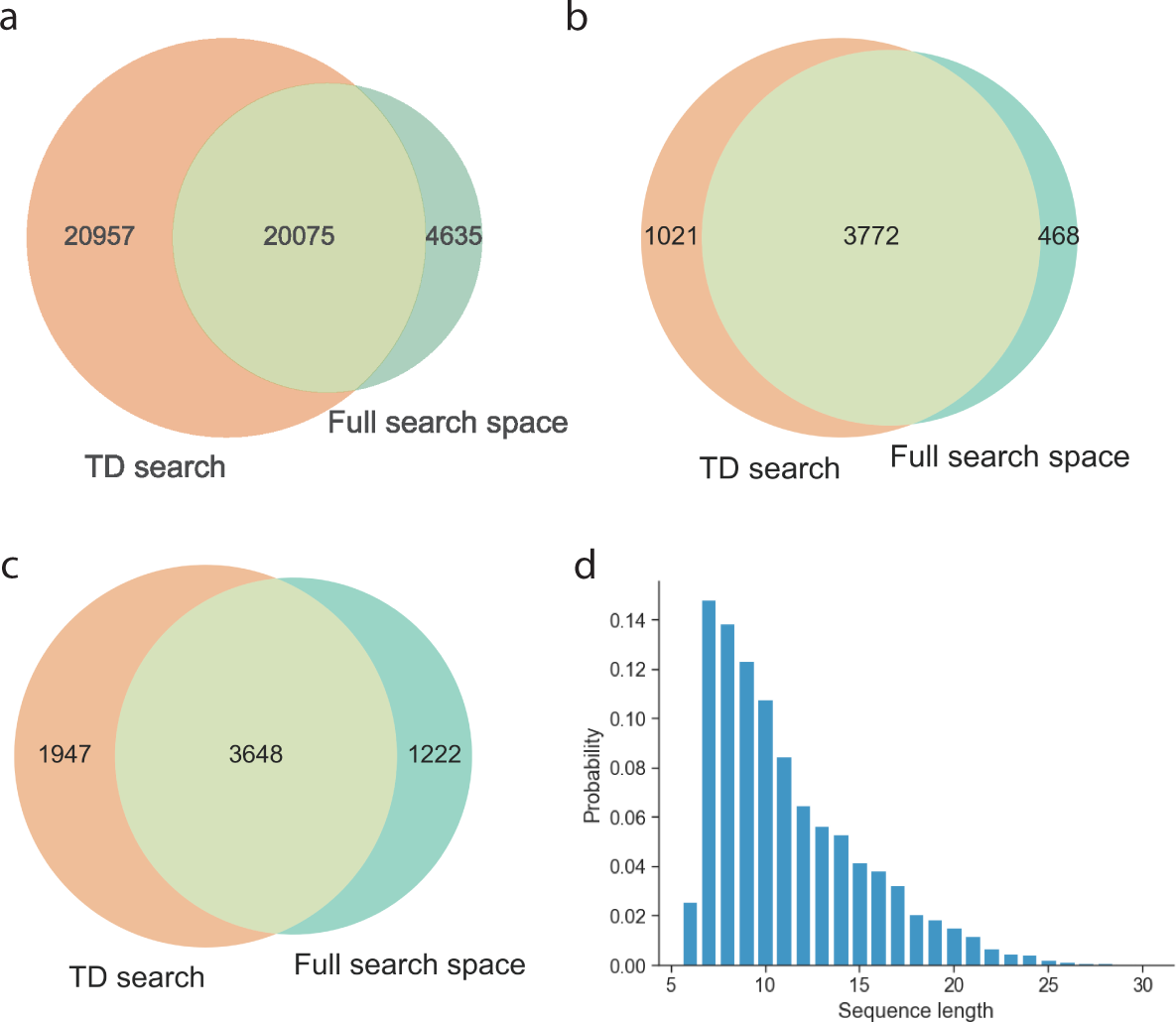
IntaNovo accurately predicts and expands detection rates in HeLa GluC degradome. **a,** Unique peptide sequences of database search and InstaNovo predicted peptides matching to the human reference proteome at 5% FDR. **b,** Proteins detected by predicted peptide sequences InstaNovo at 5% FDR. **c,** GluC candidate cleavages identified at 5% FDR (preceded by glutamate residue). **d,** Sequence length distribution for GluC generated peptides (preceded by glutamate residue).

**Supplementary Table 1.** Database search results for the datasets used in this study at 1% FDR, except for immunopeptidomics (no protein FDR)

| *Dataset* | *MS/MS* | *TD-search PSMs* | *TD-search Peptides* | *TD-search Proteins* |
| --- | --- | --- | --- | --- |
| HeLa single shot | 463,777 | 25,107 | 21,104 | 3,783 |
| Nanobodies | 257,701 | 23,147 | 5,897 | 924 |
| Herceptin | 58,609 | 1,796 | 129 | 2 |
| Wound fluids | 100,054 | 20,699 | 8,307 | 1,096 |
| *S. brodae* | 26,099 | 9,068 | 7,881 | 1,694 |
| Snake venoms | 558,247 | 21,257 | 3,446 | 610 |
| Immunopeptidome | 404,062 | 99,178 | 20,904 | 5,948 |
| HeLa degradomics | 204,831 | 115,470 | 41,483 | 4,438 |

**Supplementary Table 2.** InstaNovo evaluation results on all datasets. Confidence intervals are calculated as *±*1.96 *×* 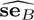 where 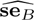 is a bootstrap standard error estimated from 10,000 replicates.*We do not calculate bootstrap standard errors for the ProteomeTools datasets because their size makes it prohibitively costly but also implies the standard errors would be very small.

| *Dataset* | AA-level performance |  |  | Peptide-level performance |  |
| --- | --- | --- | --- | --- | --- |
|  | *Error rate* | *Precision* | *Accuracy* | *Accuracy* | *AUC* |
| HeLa single-shot | 0.330 ± 0.005 | 0.609 ± 0.007 | 0.608 ± 0.007 | 0.503 ± 0.007 | 0.465 ± 0.008 |
| Immunopeptidomics | 0.211 ± 0.012 | 0.778 ± 0.020 | 0.779 ± 0.020 | 0.581 ± 0.036 | 0.532 ± 0.045 |
| *S. Brodae* | 0.204 ± 0.004 | 0.815 ± 0.008 | 0.815 ± 0.008 | 0.724 ± 0.010 | 0.697 ± 0.011 |
| Snake Venoms | 0.494 ± 0.006 | 0.396 ± 0.008 | 0.398 ± 0.008 | 0.196 ± 0.008 | 0.167 ± 0.009 |
| Nanobodies | 0.339 ± 0.004 | 0.597 ± 0.006 | 0.595 ± 0.006 | 0.447 ± 0.007 | 0.412 ± 0.007 |
| Wound Fluids | 0.467 ± 0.009 | 0.411 ± 0.012 | 0.406 ± 0.012 | 0.225 ± 0.014 | 0.190 ± 0.014 |
| HeLa degradome | 0.178 ± 0.001 | 0.798 ± 0.002 | 0.798 ± 0.002 | 0.695 ± 0.003 | 0.676 ± 0.003 |
| Herceptin | 0.215 ± 0.015 | 0.659 ± 0.029 | 0.658 ± 0.029 | 0.494 ± 0.035 | 0.472 ± 0.037 |
| Exc. Yeast | 0.371 ± 0.003 | 0.695 ± 0.005 | 0.664 ± 0.006 | 0.533 ± 0.006 | 0.497 ± 0.006 |
| HC-PT* | 0.279 | 0.687 | 0.685 | 0.573 | 0.550 |
| AC-PT* | 0.193 | 0.794 | 0.794 | 0.685 | 0.666 |
| *mean* | 0.298 | 0.658 | 0.655 | 0.514 | 0.490 |
| *std* | 0.111 | 0.147 | 0.147 | 0.174 | 0.181 |

**Supplementary Table 3.** InstaNovo+ evaluation results on all datasets. Confidence intervals are calculated as *±*1.96 *×* 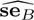 where 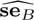 is a bootstrap standard error estimated from 10,000 replicates. *We do not calculate bootstrap standard errors for the ProteomeTools datasets because their size makes it prohibitively costly but also implies the standard errors would be very small.

| Dataset | AA-level performance |  |  | Peptide-level performance |  |
| --- | --- | --- | --- | --- | --- |
|  | Error rate | Precision | Accuracy | Accuracy | AUC |
| HeLa single-shot | 0.321 ± 0.004 | 0.617 ± 0.007 | 0.616 ± 0.007 | 0.517 ± 0.007 | 0.477 ± 0.008 |
| Immunopeptidomics | 0.161 ± 0.012 | 0.839 ± 0.022 | 0.839 ± 0.022 | 0.697 ± 0.036 | 0.644 ± 0.044 |
| *S. Brodae* | 0.187 ± 0.004 | 0.821 ± 0.008 | 0.820 ± 0.008 | 0.736 ± 0.009 | 0.697 ± 0.011 |
| Snake Venoms | 0.493 ± 0.006 | 0.393 ± 0.008 | 0.395 ± 0.008 | 0.198 ± 0.008 | 0.137 ± 0.009 |
| Nanobodies | 0.329 ± 0.004 | 0.608 ± 0.006 | 0.606 ± 0.006 | 0.464 ± 0.007 | 0.417 ± 0.008 |
| Wound Fluids | 0.467 ± 0.009 | 0.412 ± 0.012 | 0.406 ± 0.012 | 0.229 ± 0.013 | 0.166 ± 0.014 |
| HeLa degradome | 0.163 ± 0.001 | 0.811 ± 0.002 | 0.810 ± 0.002 | 0.719 ± 0.003 | 0.689 ± 0.003 |
| Herceptin | 0.192 ± 0.014 | 0.710 ± 0.028 | 0.709 ± 0.028 | 0.562 ± 0.034 | 0.526 ± 0.038 |
| Exc. Yeast | 0.244 ± 0.003 | 0.732 ± 0.005 | 0.699 ± 0.005 | 0.579 ± 0.006 | 0.556 ± 0.006 |
| HC-PT* | 0.268 | 0.696 | 0.694 | 0.589 | 0.542 |
| AC-PT* | 0.180 | 0.809 | 0.809 | 0.710 | 0.680 |
| mean | 0.269 | 0.677 | 0.673 | 0.545 | 0.503 |
| std | 0.120 | 0.157 | 0.157 | 0.187 | 0.196 |

### Algorithm 1 Knapsack chart generation by DFS

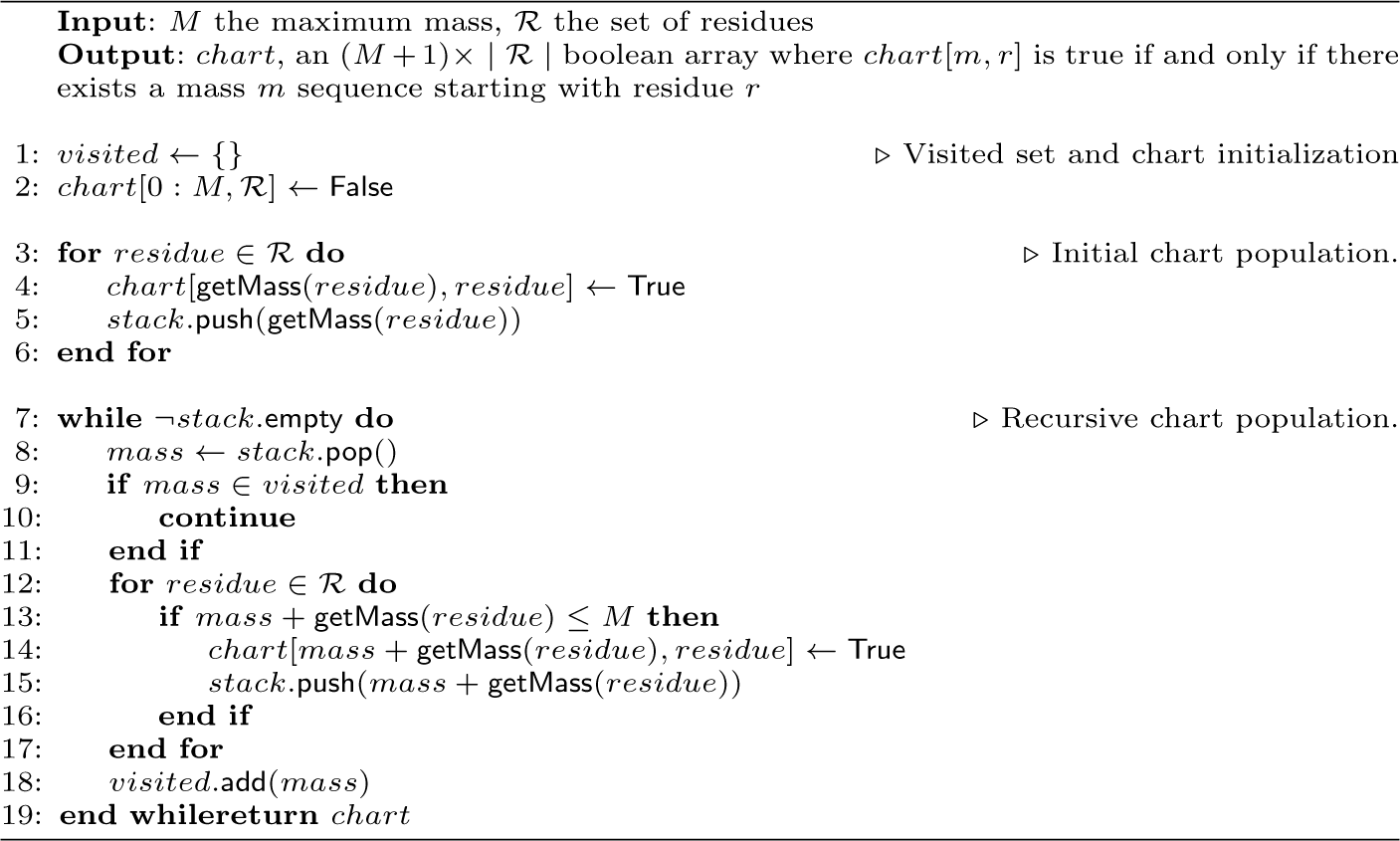

**Supplementary Table 4.** General training setup and model hyperparameters. InstaNovo+ uses the same parameters as InstaNovo unless otherwise specified.

| *Parameter* | *Description* | *Value* |
| --- | --- | --- |
| *Training hyperparameters* |  |  |
| framework | Python implementation framework | PyTorch Lightning |
| optimiser | PyTorch Lightning model optimiser | Adam |
| batch_size | Optimiser number of samples per batch | 32 |
| learning_rate | Optimiser learning rate | 5e-5 |
| epochs | Number of training epochs per experiment | 30 |
| warmup_steps | Number of LR linear warmup steps | 100,000 |
| n_tune_epochs | Maximum number of fine-tuning epochs | 6 |
| *InstaNovo hyperparameters* |  |  |
| dim_model | Hidden dimension of the model | 768 |
| n_head | Number of attention heads per layer | 16 |
| dim_feedforward | Feedforward transformer block dimension | 1,024 |
| n_layers | Number of transformer layers | 9 |
| dropout | Layer dropout probability | 0.1 |
| isotope_error | Maximum number of isotopes considered | 1 |
| n_peaks | Number of peaks pre-filtered | 200 |
| n_beams | Number of beams in beam-search decoding | 5 |
| max_charge | Maximum charge considered | 10 |
| *InstaNovo+ hyperparameters* |  |  |
| decoder_layers | Number of diffusion decoder layers | 12 |
| n_head | Number of decoder attention heads per layer | 8 |
| T | Number of diffusion timesteps | 20 |
| dropout | Layer dropout probability | 0.1 |
| *Hardware setup A* |  |  |
| gpu | Graphics processing unit | Nvidia RTX 3090ti |
| cpu | Central processing unit | Intel i5-13600k |
| vram | Available GPU virtual memory (for models) | 24GB |
| ram | Available system memory (for datasets) | 64GB |
| *Hardware setup B* |  |  |
| gpu | Graphics processing unit | Nvidia A100-80GB |
| cpu | Central processing unit | AMD EPYC 7,742<br>(32 threads used) |
| vram | Available GPU virtual memory (for models) | 80GB |
| ram | Available system memory (for datasets) | 256GB |

**Supplementary Table 5.**
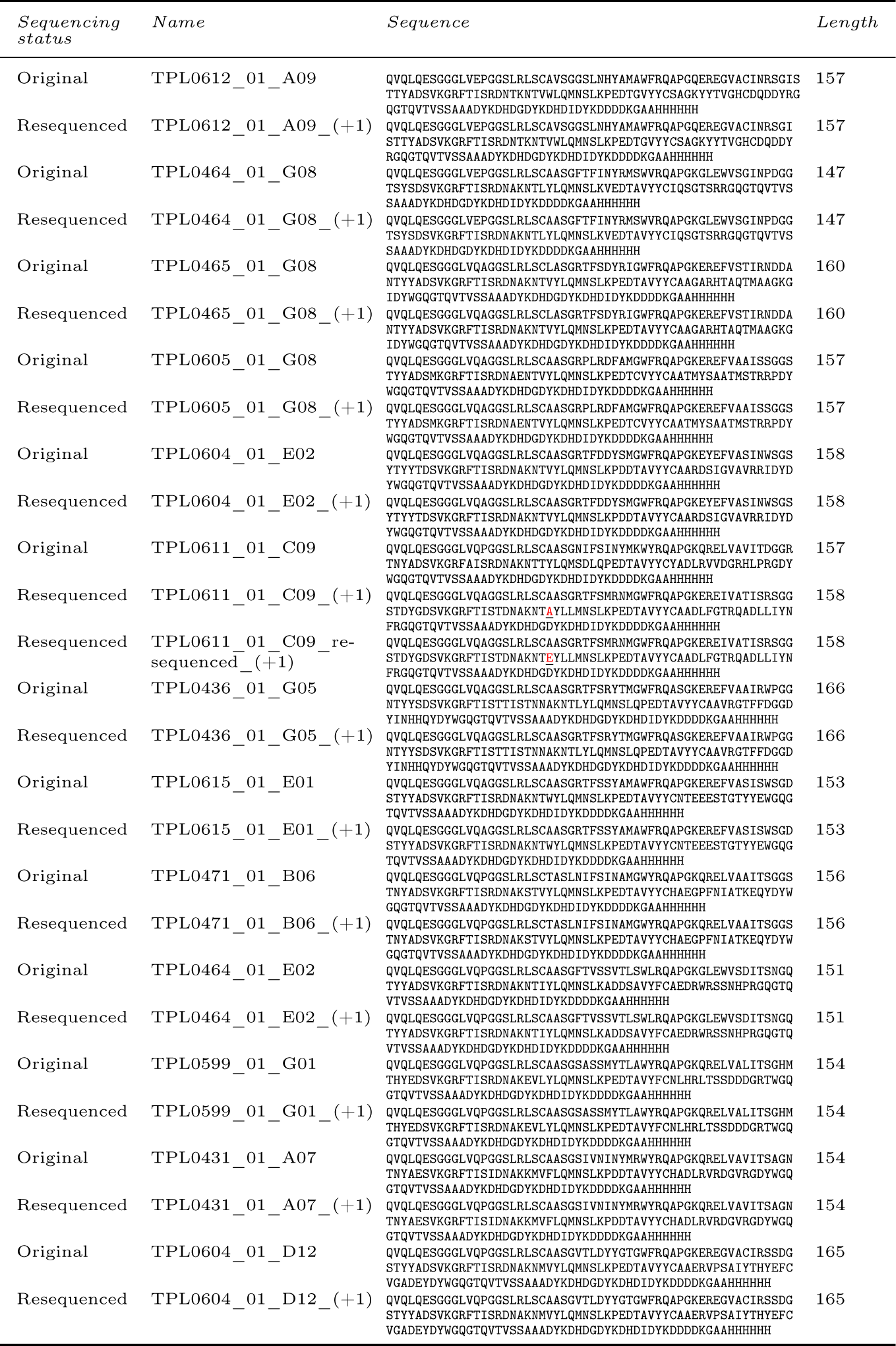
Nanobodies included in this study, including both an original Sanger sequencing round, as well as re-sequencing for sequence confirmation. Discrepancies are highlighted in red.

### Algorithm 2 Beam Search decoding with knapsack filtering

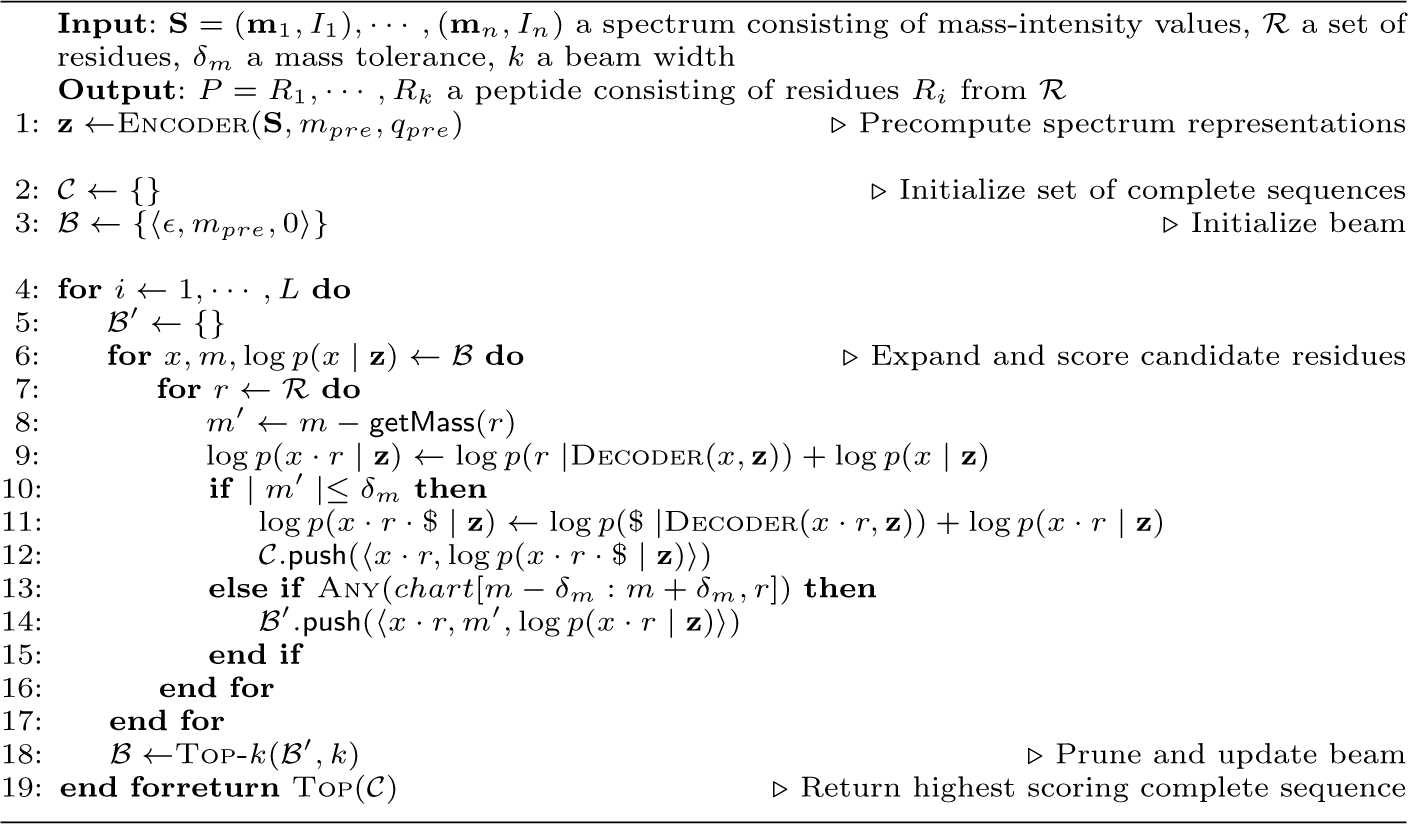

